# A new route for integron cassette dissemination among bacterial genomes

**DOI:** 10.1101/2022.09.11.507474

**Authors:** Céline Loot, Gael A. Millot, Egill Richard, Eloi Littner, Claire Vit, Frédéric Lemoine, Jean Cury, Baptiste Darracq, Théophile Niault, Delphine Lapaillerie, Vincent Parissi, Eduardo PC Rocha, Didier Mazel

## Abstract

Integrons are genetic elements involved in bacterial adaptation. They can capture, shuffle and express adaptive functions embedded in cassettes. These events are governed by the integron integrase through site-specific recombination between *attC* and *attI* integron sites. Here, we demonstrated that the integrase can efficiently catalyze insertion of cassettes in bacterial genomes, outside the *att* sites. We showed that, once inserted in genomes, cassettes can be expressed, if located near bacterial promoters, and can be excised at the insertion point and even outside, inducing chromosomal modifications in the latter case. Analysis of more than 5 × 10^5^ independent insertion events revealed a very large genomic insertion landscape with recombination sites greatly different, in terms of sequence and structure, from classical *att* sites. We named these new sites *attG*. These results unveil a new efficient route for dissemination of adaptive functions and expand the role of integrons in bacterial evolution.

## Introduction

Bacteria exchange and recombine DNA accelerating their adaptation to new stresses and environments. An active player of Gram-negative bacteria adaptation is the integron system (1). Integrons are natural genetic “toolboxes” able to stockpile, shuffle, express adaptive functions embedded in cassettes (2). Their evolutionary success relies on the diversity of these functions, among which one finds hundreds of antibiotic resistance genes. Anthropogenic pressures such as use of antibiotics have led to the selection of mobilization events in integrons, such as their association with transposons and conjugative plasmids. These so-called Mobile Integrons (MIs) have now disseminated among bacteria and constitute an important means of spreading antibiotic resistance. Importantly, MIs are only the tip of the iceberg since much larger Sedentary and Chromosomal Integrons (SCIs) have been found in approximately 17% of the available genomes in databases (3,4). Both MIs and SCIs share however the same general organization: a stable platform and a variable cassette array (Figure 1A). The platform is composed of three elements: the *intI* gene coding for a site-specific recombinase, the integron integrase, the *attI* recombination site in which promoterless cassettes are inserted, and a PC promoter oriented to direct transcription of proximal cassettes (2). The rest of the cassettes (more distal) represents a low-cost memory of valuable functions for the cell which can potentially be expressed through the reordering of the cassette array. By controlling the expression of IntI, bacteria can reshuffle the integron cassettes “on demand” in moments of stress (5). All these features make the integron a unique recombination system (5–7).

**Figure 1:**
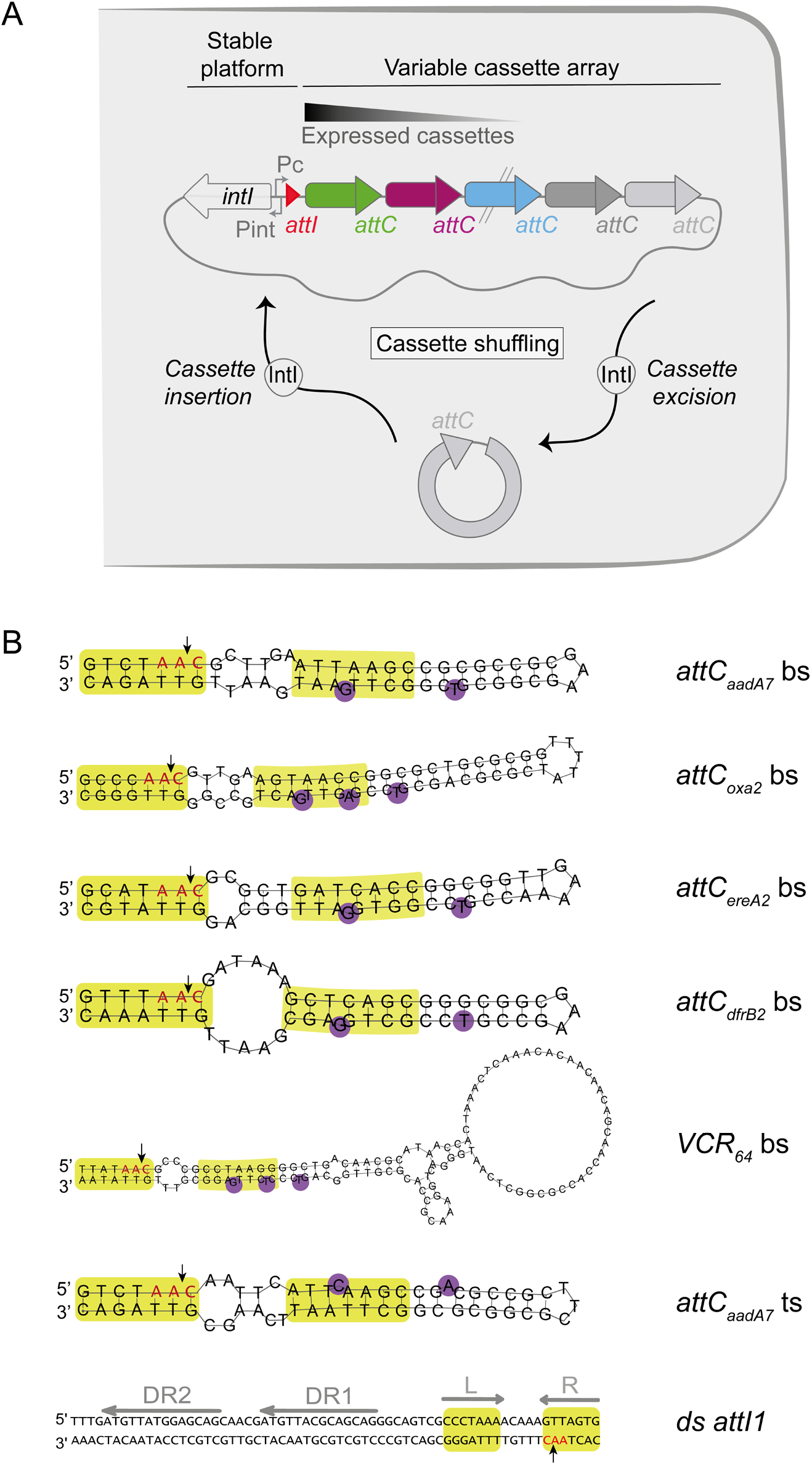
The integron system and the *att* sites. A) The Integron system The integron system is composed of the integron platform (the integrase expressing gene, *intI*, the two promoters, P_c_ and P_int_ and the *attI* recombination site (red triangle)) and of the variable cassette array. The variable cassette array contains some cassettes represented by small colored arrows. Only the first cassettes of the array are expressed, and the subsequent ones can be seen as a low-cost cassette reservoir. Upon expression of the integrase, cassette shuffling can occur through cassette excision (*attC* × *attC*) and insertion of the excised cassettes in the first position in the array (*attI* × *attC*). B) The *att* sites The sequences of the single-stranded bottom and top strands (bs and ts) of the *attC* sites and the double-stranded *attI1* (ds) site used in this study are represented. Green boxes indicate the right (R) and left (L) integrase binding sites. The 5’-AAC-3’ triplet, where the cleavage takes place, is highlighted in red and the precise cleavage point is indicated by a black arrowhead. The Direct Repeats (DR1 and DR2) of the *attI1* site are shown by grey arrows.

A cassette is a mobilizable element that generally contains a coding sequence (CDS) ended by an *attC* site. The cassette reordering is ensured by cassette excision events through recombination between two consecutive *attC* sites followed by insertion events of the excised cassettes in the *attI* site (*attC* × *attI* recombination, Figure 1A). *attC* sites share little sequence conservation and are instead recognized by the integrase through the single-stranded structures formed by their bottom strands (Figure 1B, (8)). This strand selectivity is essential to the insertion of cassettes in the correct orientation relatively to the PC promoter and hence to the expression of the promoterless gene they contain.

In MIs, many of the identified cassettes provide resistance to antibiotics, whereas cassettes found in SCIs are involved in mobility, metabolism, biofilm formation, bacteriophage resistance or host surface polysaccharide modification (2,4,9). Here, we focused on the ability of bacteria to access these adaptive functions and to build a repertoire of cassettes pertinent for their lifestyle. Our genomics analysis revealed numerous occurrences of isolated cassettes in a wide range of sequenced bacterial genomes available in databases. In line with the nomenclature proposed previously (3), we named these cassettes, SALIN, for <u>S</u>ingle *<u>a</u>ttC* site <u>l</u>acking <u>int</u>egron-integrase and we revealed the mechanism of their formation. We proposed that they may result from cassette insertion in genomes. By performing a series of *in vivo* experiments, we confirmed the high propensity of integron cassettes to disseminate into bacterial genomes. We demonstrated that these inserted cassettes can be expressed but also excised in such a way that it induces genome modifications. Deep sequencing of cassette libraries inserted *de novo* into the *E. coli* genome has shown that cassettes can target a very large number of unique sites and has allowed to characterize these insertion sites. Surprisingly, they differ greatly, in terms of sequence and structure, from both classical *attC* and *attI* recombination sites. We named these new sites, *“attG”* for <u>att</u>achment site of the genome.

These results revisit the classical model of cassette recombination and reveal a new and efficient pathway for cassette dissemination extending the role of integrons in bacterial evolution

## METHODS

### Bacterial strains, plasmids and primers

The different plasmids, strains and primers are listed in Table S1.

### Media

*Escherichia coli* and *Vibrio cholerae* were grown in Luria Bertani broth (LB) at 37°C. *E. coli* strains containing a plasmid with a thermo-sensitive origin of replication were grown at 30°C. In the case of *E. coli*, antibiotics were used at the following concentrations: carbenicillin (Carb), 100 μg/ml, kanamycin (Km), 25 μg/ml, chloramphenicol (Cm), 25 μg/ml, spectinomycin (Sp), 50 μg/ml. Diaminopimelic acid (DAP) was supplemented when necessary to a final concentration of 0.3 mM. To induce the P_bad_ promoter, L-arabinose (Ara) was added to a final concentration of 2mg/ml; to repress it, glucose (Glc) was added to a final concentration of 10mg/ml. To induce the P_tet_ promoter, anhydrotetracycline (aTc) was added to a final concentration of 1μg/ml. *V. cholerae* strains were cultivated in the same conditions and with the same antibiotic concentrations except for Cm and Sp, that were supplemented at a final concentration of 5 μg/ml and 100 μg/ml respectively. When *V. cholerae* strains were cultivated in presence of glucose, the later concentration of Sp was increased 2-fold (200 μg/ml).

### Genomics analysis of isolated inserted cassettes

#### Data

The genome dataset analyzed in the current study is the same as in (4), consisting of 21,105 complete genomes downloaded from the NCBI RefSeq database of high quality non-redundant prokaryotic genomes (accessed on 30 March 2021, https://ftp.ncbi.nlm.nih.gov/genomes/refseq/). Using IntegronFinder v2 (4), we screened the dataset to automatically and accurately identify integrons, CALINs (clusters of *attC* sites lacking integron-integrases), and isolated integrases (In0).

#### SALIN analysis in bacterial genomes

SALIN elements are single *attC* sites lacking integrase. They are a subtype of CALINs where the cluster comprises solely an *attC*. SALINs were detected with IntegronFinder v2 with the option “--calin-threshold 1” to prevent filtering out single *attC* sites.

#### Distribution of SALINs across bacterial phyla

Each genome of the dataset was assigned to a bacterial phylum according to its taxonomic identifier retrieved from the NCBI Taxonomy database. Phyla were organized in a cladogram adapted from the tree published in (10). Only phyla comprising at least one genome from our dataset were considered to build the cladogram.

### Suicide conjugation assays

The conjugation assays were based on that of Vit et al, 2021 (11). This assay mimics the natural conditions in which cassettes are delivered through horizontal gene transfer. Conjugation ensures the delivery of one of the recombination substrates (*att* sites) carried by a pSW23T suicide plasmid into a recipient cell containing a plasmid (pBAD43 or pBAD18) with or without integrase and containing, or not, a second *att* recombination site (*attI* or *attC* sites) carried on pSU38, pBAD43 or pTOPO derivate plasmids. From the donor *E. coli* ß2163 strain, the pSW23T plasmid is delivered in a single stranded form into a recipient *E. coli* MG1655 or *V. cholerae* N16961. This plasmid contains an RP4 origin of transfer (*oriT*RP4) oriented in such way to deliver either the reactive bottom strand of the *attC* recombination sites or the top one. Recipient strains cannot sustain replication of this suicide pSW plasmid and the only way for the pSW vector to be maintained in recipient cells is to recombine with *att* sites contained in the recipient strain or to insert in the bacterial genome. The frequency of cassette recombination is measured by comparing the number of recombined cells having acquired the resistance marker carried by the pSW23T vector, and the total number of recipient cells. The donor strains were grown overnight in LB media supplemented with Chloramphenicol (Cm) (resistance of the pSW plasmid), Kanamycin (Km) (resistance of the ß2163 strain) and DAP (since ß2163 donor strain requires DAP to grow in rich medium), the recipient strain was grown overnight in LB media supplemented with appropriate antibiotics depending on the plasmids used and Glucose (Glc, to repress the integrase gene when pBAD promoter is used). Both donor and recipient overnight cultures were diluted 1/100 in LB with DAP or Arabinose (Ara) respectively and incubated until OD=0.7-0.8. 1ml of each culture were then mixed and centrifuged at 6000rpm for 6mins. The pellet was suspended in 100μl LB, spread on a conjugation membrane (mixed cellulose ester membrane from Millipore, 47mm diameter and 0.45μm pore size) over a LB+agarose+DAP+Ara Petri dish and incubated 3H for conjugation and recombination to take place. The membrane with the cells was then resuspended in 5ml LB, after which serial 1:10 dilutions were made and plate on LB+agarose media supplemented with appropriate antibiotics. The recombination frequency was calculated as the ratio of recombinant CFUs, obtained on plates containing Cm and the antibiotics corresponding to recipient cells, to the total number of recipient CFUs, obtained on plates containing only antibiotics corresponding to recipient cells. The overall recombination frequency is a mean of at least 3 independent experiments.

Note that we adapted this protocol when using *V. cholerae* as recipient strain in which plasmids are more easily lost in absence of antibiotic selection than in *E. coli* (11,12).

#### Conjugation assay in presence of integron

To determine the genome cassette insertion frequency in the presence of an integron, we performed a conjugation assay proceeding exactly as described. We used the *attC_aadA7_*-containing pSW plasmid (pD060) as donor plasmid and we constructed a temperature-sensitive replicating receptor plasmid containing an *attI1* site followed by a spectinomycin resistant *attC_aadA1_* cassette (pM335) (Figure S3A). Once constructed, this vector was transformed into the MG1655 recipient strain containing the pBAD18 plasmid with or without integrase (p3938 and p979 respectively, Carb^R^). These donor and recipient strains were conjugated for 3h but at 30°C. Thus, the conjugation and recombination reactions were performed in presence of the synthetic integron *i.e*., in condition under which the plasmid can replicate (at 30°C). The membrane with the cells was then resuspended in 2ml LB+Carb+Glc, divided in two parts and incubated 24H at 30°C and 42°C. These incubation temperatures respectively favor and disfavor the pM335 plasmid maintenance. The recombination frequency of cassettes was calculated as the ratio of recombinant CFUs, obtained on Cm (to select cassette insertion), Carb (resistance of the used integrase-carrying plasmid) and Glc (to repress the *intI1* gene) containing plates, to the total number of recipient CFUs, obtained on Carb and Glc containing plates. Note that plates were incubated *in parallel* at 30°C and at 42°C. At 30°C, all cassette insertion events (in *attI1*, *attC_aadA1_* and in genome sites) are selected. At 42°C, insertion events corresponding to insertion in the recipient plasmid are counter selected and those corresponding to insertion in the chromosome are therefore enriched and more easily detectable. The overall recombination frequency is a mean of at least 3 independent experiments.

#### Cassette expression assay

To test if *attC* cassettes can be expressed once inserted in genome, we performed a conjugation assay proceeding exactly as described above. We constructed a suicide plasmid vector containing the *attC_aadA7_* but adding a kanamycin (*km*) resistance gene without promoter just downstream the *attC* site (pN695-pN697, Cm^R^). We also added two different RBS just upstream the *km* gene (RBS1=GGAGG, pN708-pN709 and RBS2=AGGAG, pN705-pN707, Cm^R^). Once constructed, these plasmids were transformed into the β2163 donor strain. These donor strains and the recipient MG1655 strain containing the pBAD43 plasmid with or without integrase (pL294 and pL290 respectively, Sp^R^) were conjugated. The recombination frequency of cassettes was calculated as the ratio of recombinant CFUs, obtained on Cm (to select cassette insertion), Sp (resistance of the pBAD43 plasmid) and Glc (to repress the *intI1* gene) containing plates, to the total number of recipient CFUs, obtained on Sp and Glc containing plates. The recombination frequency of solely cassettes expressing the *km* resistance gene was calculated as the ratio of recombinant CFUs, obtained on Km, Cm, Sp and Glc containing plates, to the total number of recipient CFUs, obtained on Sp and Glc containing plates. The overall recombination frequency is a mean of at least 3 independent experiments.

#### Cassette excision assay

To test if *attC* cassettes can be excised once inserted in genome, we performed a conjugation assay proceeding exactly as described above. We constructed a new suicide plasmid vector containing the *attC_aadA7_* and the *ccdB* toxin gene under the control of the PBAD promoter (pM779-pM781, Cm^R^). The *ccdB* gene was previously used as a potent counterselection marker in several commonly used applications (13,14). Once constructed, this plasmid was transformed into the β3914 donor strain to perform conjugation (a *pir+* CcdB resistant *E. coli* strain (13)). We also constructed a new vector expressing the integrase. This vector is a pBAD43 temperature-sensitive replicating vector with a P_TET_ promoter in place of the previously used P_BAD_ promoter (pN435-pN440, Km^R^). Once constructed, this plasmid was transformed into the *MG1655ΔrecA* recipient strain. Both donor and recipient strains were conjugated. Conjugation was performed by adding anhydrotetracycline (aTc) in place of Ara to express the integrase gene and at 30°C. We also used Glc to repress the *ccdB* toxin gene. Cassette insertion events in genome were selected by plating cells on Cm (to select cassette insertion) and Glc (to repress the *ccdB* toxin gene) containing plates. After that, 24 recombinants clones were randomly picked and cultivated at 42°C to ensure the loss of the thermosensitive pBAD43 plasmid. The 24 obtained recombinant clones were transformed with the pBAD43::P_TET_-*intI1* plasmid (pM888, Sp^R^) and, as control, with the pBAD43::P_TET_ plasmid (pM889, Sp^R^). Clones were grown for 8h in presence of aTc (to express the integrase gene), Sp and Glc. The aim of this step is to promote successful recombination event leading to cassette excision. The excision frequency of cassettes was calculated as the ratio of recombinant CFUs, obtained on Sp and Ara containing plates, to the total number of recipient CFUs, obtained on Sp and Glc containing plates. Note that in presence of Ara (to express the *ccdB* toxin gene), only clones that have lost the *ccdB* gene, due to a cassette excision event, are selected while the others die. We checked the Cm sensitivity of a large number of recombinant clones. The overall recombination frequency is a mean of at least 3 independent experiments.

#### Testing the hotspots as receptor and donor sites

To test if the hotspots (HS), the median spot (MS) and the unique spot (US) can be used as donor or receptor sites, we performed conjugation assays proceeding exactly as described above.

To test the HS, MS and US sites as receptor sites, we used the *attC_aadA7_*-containing pSW plasmid (pD060) as donor plasmid and we constructed receptor plasmids containing the different HS, MS and US. Once constructed, each vector was transformed into the MG1655 recipient strain containing the pBAD43 with or without integrase (pL294 and pL290 respectively, Sp^R^). The donor strain and these recipient MG1655 strains were conjugated. The recombination frequency of cassettes was calculated as the ratio of recombinant CFUs, obtained on Cm (to select cassette insertion), Sp (resistance of the pBAD43 plasmid) and Glc (to repress the *intI1* gene) containing plates, to the total number of recipient CFUs, obtained on Sp and Glc containing plates. The overall recombination frequency is a mean of at least 3 independent experiments.

To test the HS as donor sites, we constructed suicide plasmid vectors containing the *ybhO* hotspot site delivering either the bottom strand (pO323-pO324) or the top one (pO321-pO322), the *alsB* hotspot site delivering either the bottom strand (pO749-pO750) or the top one (pO751) and the *pyrE* hotspot site delivering either the bottom strand (pO752) or the top one (pO753-pO755). Once constructed, these plasmids were transformed into the β2163 donor strain. These donor strains and the recipient MG1655 *recA* strain containing the pBAD43 plasmid with or without integrase (pL294 and pL290 respectively, Sp^R^) and the pSU38Δ::*attC_aadA7_* plasmid (pO371-pO372) were conjugated. The recombination frequency of cassettes was calculated as the ratio of recombinant CFUs, obtained on Cm (to select cassette insertion), Sp (resistance of the pBAD43 plasmid), Km (resistance of the pSU38 plasmid) and Glc (to repress the *intI1* gene) containing plates, to the total number of recipient CFUs, obtained on Sp, Km and Glc containing plates. The overall recombination frequency is a mean of at least 3 independent experiments.

### Analysis of recombination events and point localization

For each experiment, clones were randomly picked and isolated on antibiotic containing plates. Recombination events were checked by Polymerase chain reaction (PCR) using the DreamTaq DNA polymerase (Fisher Scientific). All the PCRs were directly performed on, at least, eight randomly chosen bacterial clones per experiment. Some PCR reactions were purified using the PCR purification kit (Fisher Scientific) and sequenced to confirm the insertion point (Eurofins).

#### Insertion events in att sites carried on pSU38 vector

For analysis of co-integrates formation, we performed PCR reactions on randomly chosen clones per each experiment using SWend/MRV primers to confirm the bs recombination (when bs is injected), SWend/MFD primers to confirm the bs recombination (when ts is injected) and Swend/MRV primers to confirm the ts recombination (when the ts is injected). Recombination points were precisely determined by sequencing PCR products using MRV or MFD primers.

#### Insertion events in HS, MS and US sites carried on pTOPO vector

For analysis of co-integrates formation, we performed PCR reactions on randomly chosen clones per each experiment using SWbeg/MFD primers to confirm the *ybhO, alsB, ilvD, pyrE* and *yjhH* hotspot recombination and using SWbeg/MRV primers to confirm the *metC* hotspot, MS-*abgA* and *US-ygcE* recombination. Recombination points were precisely determined by sequencing PCR products using Swbeg primers.

#### Insertion events in att sites carried on pSC101ts vector

For analysis of co-integrates formation, we performed PCR reactions on randomly chosen clones per each experiment using SWbeg/o1714 primers to confirm the *attI1* recombination and using SWend/o1704 primers to confirm the *attC_aadA1_* recombination. Recombination points were precisely determined by sequencing PCR products using o1714 or o1704 primers.

#### Insertion events on bacterial genome

For analysis of insertions of *attC* containing-pSW23T plasmids in *E. coli* genome, we performed random PCR (Figure S1). For these, we performed a first random PCR reaction using the o1863 degenerated and the o2405 primers. The o2405 primer hybridizes upstream of the *attC* sites on pSW23T plasmids. Due to the presence of degenerate nucleotides in the o1863 primer, low hybridization temperatures were used, first, 30°C during 5 cycles and after, 40°C during 30 cycles. The obtained amplified DNA fragments were subjected to a second PCR reaction to enrich for PCR products corresponding to cassette insertion. For this purpose, we used o1865 and o1388 primers. These primers hybridize respectively to the fixed part of the degenerated o1863 primer and upstream (but closer than o2405) of the *attC* sites on pSW23T plasmids. Recombination points were precisely determined by sequencing PCR products using o1366. The o1366 primer hybridizes upstream (but closer than o1388) of the *attC* sites on pSW23T plasmids.

We also used the same procedure to detect insertions of *attC* containing-pSW23T plasmids in the *V. cholerae* genome. Interestingly, *V. cholerae* contains a massive chromosomal integron located on the chromosome 2 and harboring 179 *attC* containing cassettes (15). However, performing random PCR on 48 randomly chosen clones and sequencing 15 of them, we did not detect any insertion events in the *attC* sites carried by the resident SCI, showing that integron cassettes can be inserted in the *V. cholerae* genome at very high frequency even in the presence of a resident integron.

#### Excision events on bacterial genome

For analysis of excision events from insertions of *attC* containing-pSW23T plasmids in *E. coli* genome, we performed PCR. Note that to perform PCR, we designed appropriate primers based on the knowledge of cassette recombination points during insertion. We performed PCR on clone 7, 8 and 9. We used o6263/o6264 for clone 7, o6265/o6266 for clone 8 and o6267/o6268 for clone 9. Excision points were precisely confirmed by sequencing the PCR products.

### Principles of insertion profiling by deep sequencing

#### Library preparation

Clones obtained from three independent conjugation assays were collected and subjected to genomic extraction using the kit DNeasy® Tissue Kit (Qiagen). DNA was mechanically fragmented using the Covaris method (DNA shearing with sonication). Adaptors are ligated to the fragmented DNA pieces. Nested PCR, including 30 rounds of amplification, was performed to amplify low-abundance junctions in the DNA population using first o1366 and o6036 (to amplify the cassette genome junction, Table S1) and second o6035 and o6036 (to reconstitute adaptors, Table S1). PCR-enriched junctions were deep-sequenced using next-generation sequencing (NGS) technologies (Illumina MiSeq v3 single-end 150 cycles) (Figure S5).

#### Bioinformatics analysis

The Nextflow pipeline used to analyze the raw fastq files and generate the Figure 4 and S6 is available at https://gitlab.pasteur.fr/gmillot/14985_loot and is briefly described here. First, non-genomic sequences such as barcodes and linkers were trimmed from the raw fastq reads. Then, reads showing the *attC* sequence in 5’, expected by the PCR-enriched junctions described above, were selected and the *attC* sequence was trimmed such that the 5’ end of reads corresponds to the insertion site in the genome. Only reads showing at least 25 nucleotides post-trimming were kept and aligned on the *E. coli* str. K-12 substr. MG1655 reference genome sequence (NCBI NC_000913.3) using the --very-sensitive option of Bowtie2 (16). Q20 mapped reads were selected and checked for absence of soft clipping in each extremity. In our experimental design, reads showing the same plasmid insertion site result from three non-exclusive processes: 1) same plasmid insertion site in two different bacteria, 2) bacteria clonal amplification and 3) DNA PCR enrichment. The MarkDuplicates-Picard tool of GATK (https://github.com/broadinstitute/gatk) could not be used to remove read duplicates, as eliminating read showing identical 5’ end would also remove reads resulting from the first process. To alleviate such stringency, we considered as duplicates reads showing the same 5’ and 3’ extremities, and we analyzed libraries with and without removing the duplicates. Position of plasmid insertion was defined by the 5’ extremity of forward and 3’ extremity of reverse read alignments. Sequence around insertion sites were extracted using bedtools (https://bedtools.readthedocs.io/en/latest/) and the nucleotide consensus of these n sequences were visualized with the R package ggseqlogo (17). Random insertions were determined by deducing a sequence motif from the insertion consensus and by randomly select with replacement n positions among all the motif positions present in the reference genome. For instance, for the IntI1 library, the obtained GWT consensus motif has been found present 338,348 times in the *E. coli* reference genome: 169,209 and 169,139 times in the top and bottom strand, respectively. From here, we randomly selected with replacement 361,464 sites (number of insertion sites after read duplicate removal) among the 338,348 5’GWT3’ sites present in the genome to use them as a “random control” of cassette insertion (Figure 4 and S6).

#### Digital PCR

The absolute quantification of *ori* and *ter* was performed in multiplex by digital PCR (Stilla Technologies, Villejuif, France) and used to generate the *oriC/terC* ratio. Digital PCR was based on that of Madic et al, 2016 and de Lemos et al, 2021 (18,19). Genomic DNA was purified from the beginning of the conjugation to 60 min, the time frame during which most conjugation events occur. The primers and probes used are listed in Table S1. The *oriC/terC* ratio was found to be ~2, indicating that cells contain twice as many *oriC* than *terC* copies.

## RESULTS

### *In silico* detection of isolated cassettes in bacterial genomes

We took advantage of the recent release of IntegronFinder 2.0 (4) to search for isolated cassettes in the 21,105 complete bacterial genomes retrieved from NCBI RefSeq database. We defined isolated cassettes as CDS associated with an *attC* site with no other *att* integration sites and integrase gene nearby. We found 2469 genomes containing isolated cassettes (Figure 2A). By analogy with the previously described CALINs (<u>C</u>lusters of *<u>a</u>ttC* sites lacking integron-integrases, (3)), we called these isolated cassettes SALINs (<u>S</u>ingle *<u>a</u>ttC* site lacking integron-integrases). Among the 2469 genomes containing SALINs, 1847 contain only one SALIN and the remaining 622 more than one SALIN (Figure 2B). Interestingly, SALINs are more represented than CALINs (3400 SALINs versus 2961 CALINs, Figure 2C). CALINs are thought to arise from integrons by integrase gene loss caused by deletions or pseudogenization events or by rearrangements of parts of the cassette array mediated by transposable elements (3). However, previous analysis showed that most CALINs (95%) are not close to recognizable *intI* pseudogenes (3). Furthermore, we observe that many CALINs are not surrounded by transposable elements and tend to be small: among the 2961 CALINs, 1362 harbor only 2 cassettes (Figure 2C). Interestingly, we found certain phyla such as the Tenericutes in which we only observed SALINs (Figure 2D). This raises the possibility that, rather than being remnants of integrons, some of small CALINs and SALINs could result from the insertion of integron cassettes in bacterial genomes.

**Figure 2:**
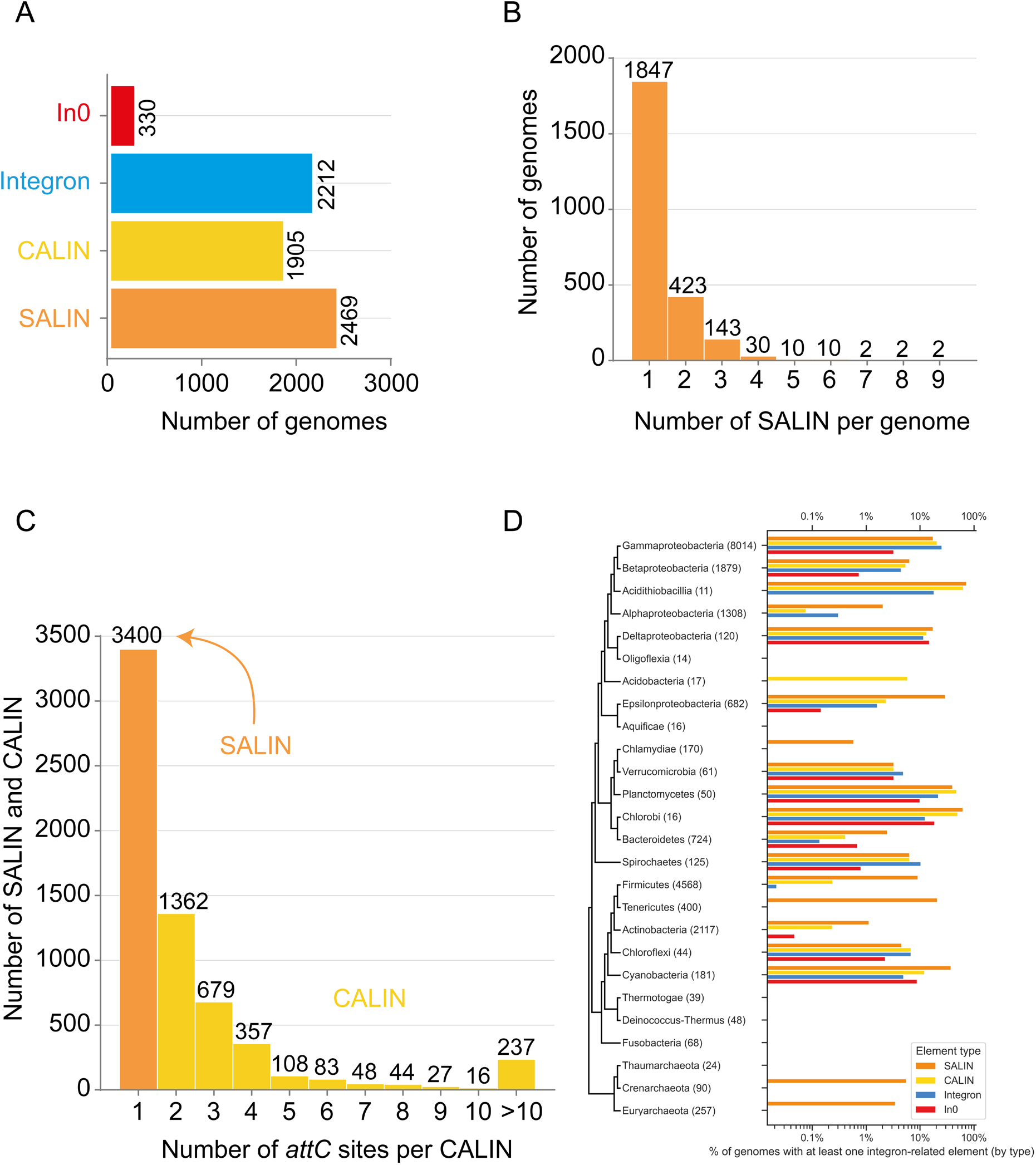
Distribution of integrons across bacteria using the RefSeq NCBI database. A) Number of bacterial genomes containing either a complete integron (integron), an integrase gene (In0), a Cluster of *attC* sites lacking integron-integrase (CALIN) or a Single *attC* site lacking integron-integrase (SALIN) B) Number of SALIN found per genome C) Number of *attC* sites per CALIN D) Taxonomic distribution of SALINs, CALINs, In0 and Integrons across major bacteria phyla

### Integron cassettes can disseminate in bacterial genomes

To validate our hypothesis, we tested the insertion capability of integron cassettes in bacterial genomes. Using our suicide conjugation assay, we delivered *attC-containing* plasmids (mimicking cassettes) on a single-stranded form in recipient strains containing a vector expressing or not the integrase (Figure 3A) (8,11,20). We tested the insertion properties of different *attC* sites from both MIs and SCIs (*attC_aadA7_, attC_oxa2_, attC_ereA2_, attC_dfrB2_* or VCR sites) chosen for their high recombinogenic properties (Figure 1B, (21)). All these sites showed a rate of recombination comprised between 10^-2^ and 10^-4^ (Figure 3A), far above what was obtained without integrase expression. Using random PCR and sequencing approaches (Figure S1 and Methods), we confirmed that cassettes were inserted at several locations in the *E. coli* genome. Furthermore, the rate of insertion dropped to 10^-6^ when using the *attI1* site or the top strand of the *attC_aadA7_* site, equivalent to the rates obtained in the absence of IntI1. Thus, *attI* sites do not recombine in genomes at a significant extent, whereas *attC* sites can recombine in genomes in the same way that they recombine with the *attI* and *attC* sites, *i.e*., as a single-stranded structured form made by the bottom strand, but not by the top one (8).

**Figure 3:**
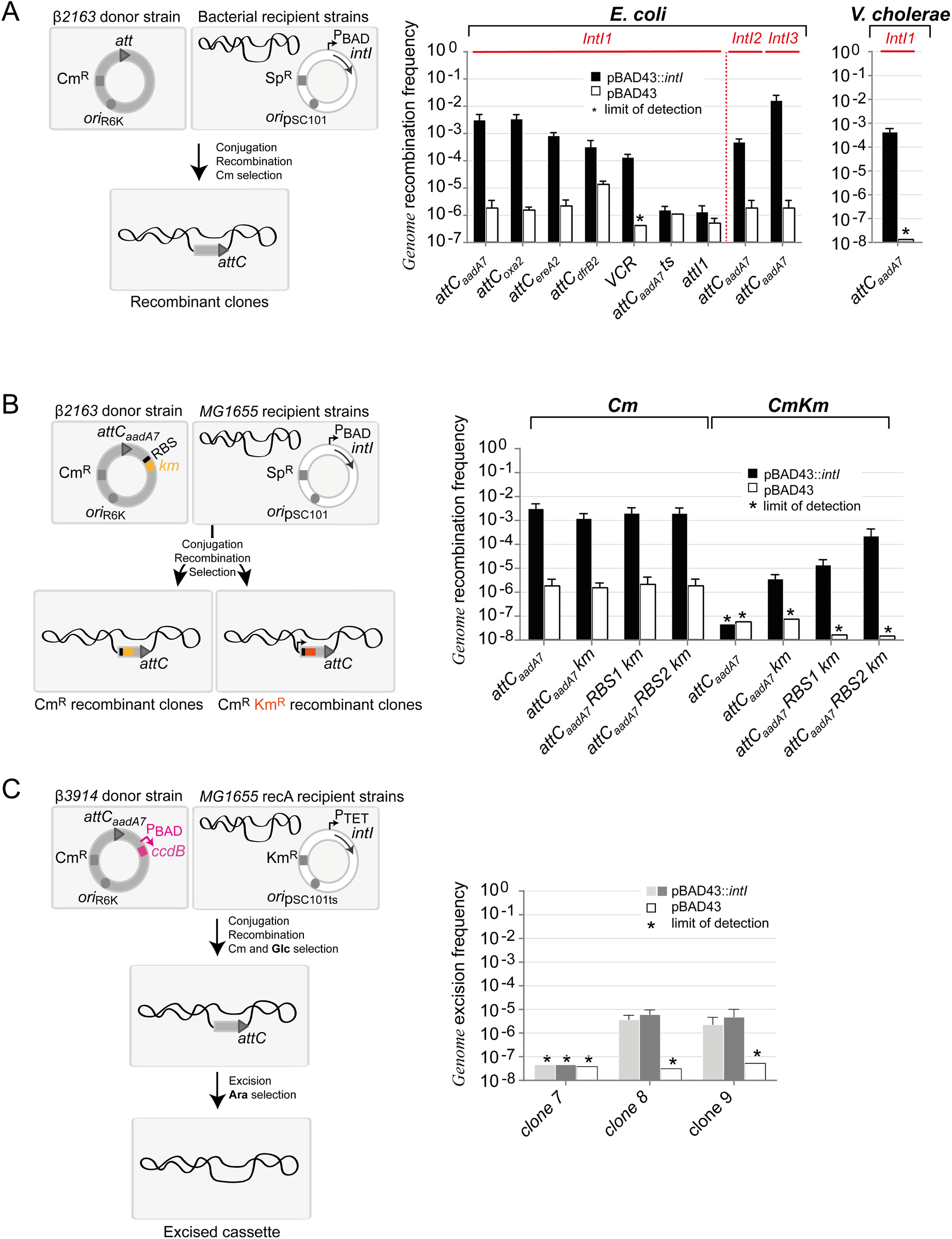
Cassette insertion in genome sites. A) Cassette insertion in *Escherichia coli* and *Vibrio cholerae* genome sites The experimental setup of the cassette insertion assay is described (left panel). The pSW23T suicide vectors containing the *attC* sites are delivered by conjugation from the ²2163 *E. coli* donor strains to the MG1655 *E. coli* or N16961 *V. cholerae* recipient strains. The *attC* sites carried by the suicide vectors are represented by a grey triangle. The recipient strains contain a plasmid expressing, or not, the integron integrase (*intI*). The inducible PBAD promoter is represented by a black arrow. Phenotypic resistances to chloramphenicol (Cm^R^) and spectinomycin (Sp^R^) are represented by grey rectangles and origin of replication by grey circles. As pSW23T cannot replicate in the recipient strains, recombinant clones can be selected on appropriate Cm containing plates to evaluate the *genome* recombination frequency (right panel, see Methods). The graph representing all the recombination frequencies obtained in several conditions is shown (Right panel). The donor plasmids are indicated in the axis-x legends (see also Table S1). The recipient strains (*E. coli* or *V. cholerae*) and the expressed integrases (IntI1, IntI2 or IntI3) are indicated at the top of the bars. pBAD43::IntI and pBAD43 means that recipient strains contain the pBAD43 integrase expressing vector or the empty pBAD43 vector, respectively. Asterisk (*) indicates the recombination frequency was below detection level, indicated by the bar height (limit of detection). Values represent the mean of at least three independent experiments and error bars represent mean absolute error. ts, means that top strand is injected during conjugation. B) Expression of inserted cassettes in *Escherichia coli* genome sites The experimental setup of the cassette expression assay is described (left panel). The setup is the same as in Figure 3A, except that the donor plasmids contain an *attC_aadA7_* site and a promoterless *kamamycin* gene (orange rectangle) associated or not with a Ribosome Binding Site (RBS1 or RBS2) and that the MG1655 *E. coli* receptor strain express, or not, the IntI1 integrase. All recombinants are selected on Cm containing plates and recombinants clones expressing the *kanamycin* gene (orange rectangle) are selected on Cm and Km containing plates. A bacterial promoter is shown by a black arrow. The graph representing all the recombination frequencies obtained in several conditions is shown (Right panel). The donor plasmids are indicated in the axis-x legends (see also Table S1). C) Excision of inserted cassettes in *Escherichia coli* genome sites The experimental setup of the cassette excision assay is described (left panel). The setup is the same as in Figure 3A, except that the donor plasmid contains an *attC_aadA7_* site and the *ccdB* toxic gene (pink rectangle) under the control of the PBAD promoter (pink arrow), induced by arabinose (Ara) and repressed by glucose (Glc), and that the MG1655 *recA E. coli* receptor strain contain a thermosensitive (ts) plasmid expressing, or not, the IntI1 integrase and encoding for the kanamycin resistance (Km^R^). The integrase gene is under the control of the PTET promoter (back arrow). 24 recombinants clones were randomly selected and placed at 42°C to remove the thermosensitive plasmid and re-transformed with plasmid expressing or not the IntI1 integrase. Recombinants clones corresponding to excised cassettes are selected on arabinose (Ara) containing plates. The graph representing the recombination frequencies obtained for three clones (7, 8 and 9) is shown (Right panel). Excision frequencies at the insertion point (dark grey bars) and outside the insertion point (light grey bars) are indicated for each clone.

Replacing the integrase IntI1 by IntI2 or IntI3, respectively the integrases of the class 2 and class 3 MIs, still provides a high rate of insertion of the *attC_aadA7_* cassette, either in the *E. coli* genome (Figure 3A) or in a recipient plasmid carrying the *attI2/attI3* or *attC_ereA2_* canonical sites (Figure S2), indicating that the property to insert integron cassettes in the *E. coli* genome is not restricted to IntI1. We also demonstrated that IntI1 can mediate cassette insertions at high frequency (almost 10^-3^) in the genome of another Gram-negative bacteria, the pathogenic *V. cholerae* strain (Figure 3A), and this, despite the presence of the *V. cholerae* endogenous chromosomal integron (see methods, (15)).

### Genome inserted cassettes can be expressed

To determine if the promoterless integron cassettes can be expressed when inserted in the genome, we added a *kanamycin* resistance gene without promoter preceded or not by a ribosome binding site (RBS) into the donor plasmid containing the *attC_aadA7_* site (Figure 3B). This matches previous observations where some CDS in cassettes are preceded by a suitably spaced RBS (2). We tested two different RBS1 and RBS2 sites naturally found in some integron cassettes. For all tested cassettes, as expected, a high frequency (more than 10^-3^) of genomic insertions was observed in Cm selective medium (Figure 3B). Using selective medium containing Cm and Km, the insertion and expression rate was high, as soon as the donor plasmid presents the *km* resistance gene, and especially when it is preceded by a RBS motif (up to 10^-4^). Comparing the rates obtained in presence of Cm (2 ×10^-3^) and of, Cm and Km (2 ×10^-4^) indicated that up to 10% of inserted cassettes can be expressed when carrying the RBS2. Performing random PCR and sequencing on over 40 randomly chosen Km^R^ clones confirmed that the expressed cassettes were inserted near a resident promoter. These results demonstrate that a large proportion of genome inserted cassettes can be expressed, if they are located in the vicinity of a promoter, thus conferring a new phenotype on the bacteria.

### Genome inserted cassettes can be excised

To test if genome inserted cassettes can be excised, we added, in the *attC_aadA7_*-containing donor plasmid, a *ccdB* gene encoding a bacterial toxin under control of the P_BAD_ promoter (Figure 3C) (13,14). First, we performed our conjugation assay and selected the cassette insertion events adding glucose to repress the *ccdB* gene. Second, we randomly chose 24 recombinant clones and performed an excision assay (Methods). We selected excised clones on arabinose containing plates. In these conditions, the CcdB toxin is expressed and only clones that have lost the *ccdB* gene, due to a cassette excision event, are selected while the others die. Excision events were notable for clones 8 and 9 (Figure 3C). PCR and sequencing analysis revealed that excision events can occur at the precise insertion site or at other nearby genomic sites, inducing genome modifications for the latter case (Figure 3C and S4).

### Cassette insertion occurs in a broad number of genomic locations

Deciphering where and how integron cassettes are inserted in genomes is fundamental to understand their potential cost and impact on host evolution. We therefore performed a genome wide NGS mapping of insertion sites (Figure S5) using a library of around 50,000 recombinant clones obtained after insertion of the *attC_aadA7_* cassettes in the *E. coli* genome catalyzed by the IntI1 integrase. Genomic sequences flanking the insertions were extracted from each sequencing read, aligned to the *MG1655 E. coli* reference genome, and used to call precise insertion sites. Genomic insertion occurred in many positions in the *E. coli* genome (22,271 unique insertion sites, Figure S6), with a huge variation in site usage (Figure 4A-B). To better analyze these data, we first removed the duplicated reads from the 3,205,043 reads leading to 361,464 reads (Figure 4C-H and Figure S6). A window size of 200,000 bps sliding every 100 bps along the genome (Figure 4C) uncovered preferential insertions near the origin of replication. Performing multiplex digital PCR, we demonstrated that, during the assay, the cells contain twice as many *oriC* than *terC* copies (Figure S7) explaining, at least in part, why insertions are favored near the origin of replication. Alignment of all the DNA sequences flanking the integration cutting sites revealed a short 5’GWT3’ consensus sequence (Figure 4D). No obvious bias of insertion was detected depending on the forward or reverse strand of the genome (Figure 4E, left panel), and the same for the leading or lagging strand template during replication (Figure 4E, right panel). As expected, a decrease of insertions in the essential genes of the *E. coli* genes (7% (295/4213) according to (22)), when compared to non-essential genes or when using random insertions (Figure 4F-G) was detected, but with no obvious effect of insertion in the same or opposite direction of transcription (Figure 7G). Finally, insertion around transcription start sites (TSS) appeared shifted upstream of essential genes (-105 bps, Figure 7H). Altogether, we conclude that cassette insertion can occur all along the *E. coli* genome with no notable effect of genomic features, except the 5’GWT3’ DNA sequence motif and the avoidance of essential genes. Interestingly, same results were observed using IntI2 and IntI3 (Figure S6). The only remarkable difference was a 1 base larger 5’TGWT3’ consensus insertion site for IntI2 that may explain the lower frequency of cassette insertion in genomes mediated by this integrase (Figure 3A).

**Figure 4:**
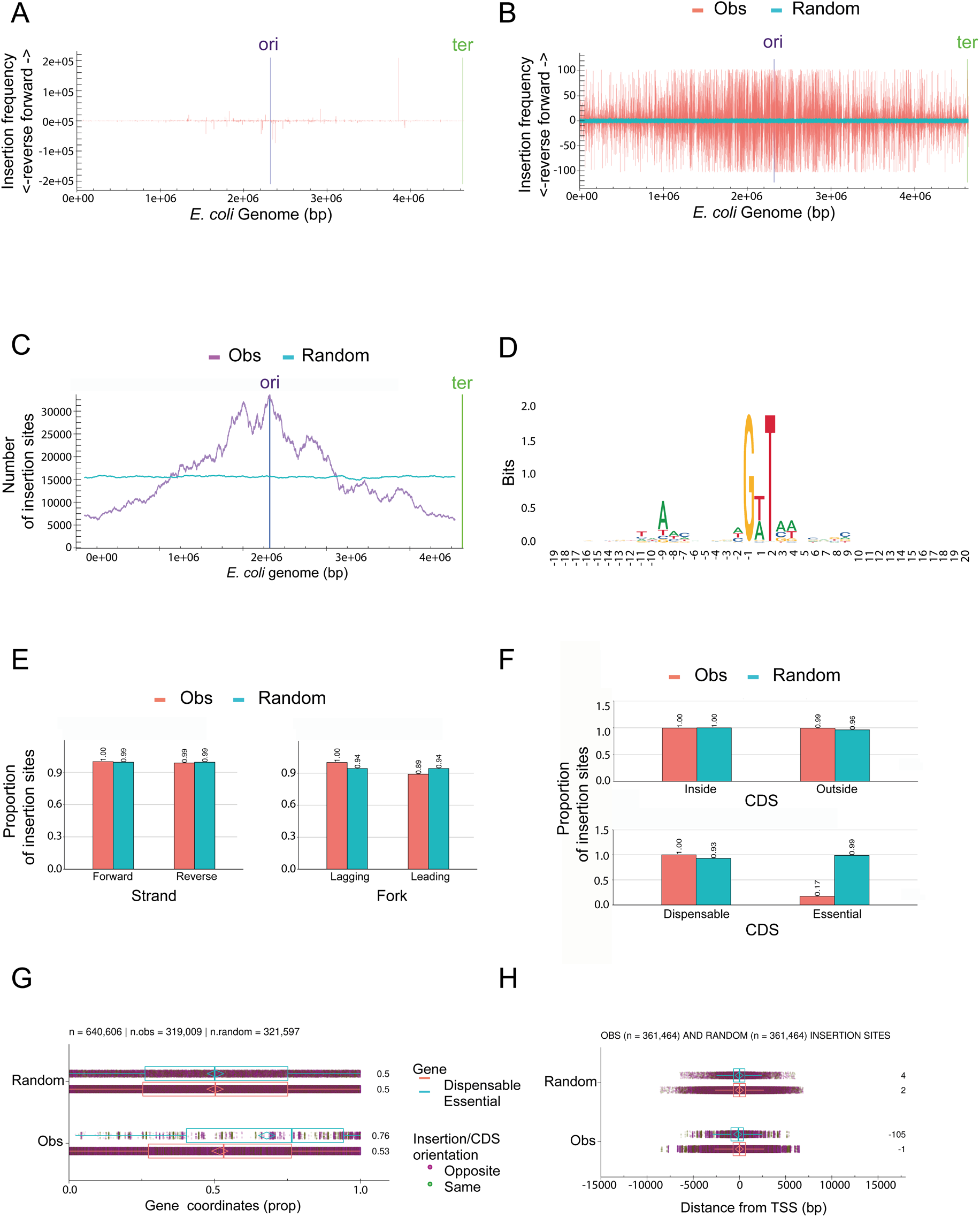
Computational analysis of Deep sequencing data. A) Insertion site usage all along the *E. coli* genome without read duplicate removal (n = 3,205,043). Insertion in the forward or reverse strand determines the orientation of the inserted cassettes (see methods for details). bp, base pair. Ori and ter, region of origin of replication and of termination respectively. B) Insertion site usage all along the *E. coli* genome with read duplicate removal (n = 361,464) All the other panels from this figure derive from read duplicate removal. Obs, observed insertions; Random, random insertions using the GWT consensus motif of insertion (see D). C) Number of insertion sites all along the *E. coli* genome using a sliding window of 200 kb sliding every 100 bases. D) Consensus sequence of insertion sites. A total of 20 bases around the cleavage point of each read was aligned to generate the motif. The cleavage occurs between the -1 and 1 bases. Bits refers to the information content. E) Proportion of insertion sites according to the strand polarity (Forward or Reverse) and according to replication orientation. Leading and lagging mean that the read corresponds to the neo-synthetized leading and lagging strand during replication, respectively. The proportions are relative to the maximal proportion set to 1. F) Proportion of insertion sites all along the *E. coli* genome according to replication orientation using a sliding window of 200 kb sliding every 100 bases G) Proportion of insertion sites according to Coding sequence (CDS) in the genome, either inside/outside CDS or dispensable/essential CDS The proportions are relative to the proportion of CDS regions in the genome (8%) and relative to the maximal proportion set to 1. H) Relative position of insertion sites inside dispensable (red) or essential (blue) CDS, between 0 (start codon) and 1 (stop codon) Box, inside vertical bar, whisker and diamond indicate quartiles, median, 1.5 x Inter Quartile Range and mean, respectively. Numbers on the right side correspond to median values. Each dot represents a single insertion site in purple or gold, depending on the opposite or same plasmid orientation versus CDS orientation respectively. I) As in H but for the distance of insertion site from the closest Transcription Start Site (TSS) in base pairs (bp)

### Cassette insertions occur in hotspots in the *Escherichia coli* genome

Several insertion hotspots were denoted when considering all the reads (*i.e*., including the duplicate ones), whatever the IntI1, IntI2 or IntI3 integrase tested (Figure 4A and Figure S6). Note that for IntI1 and IntI3, the strongest hotspot was the same, located in the *ybhO* gene (Figure S6, red boxes). However, duplicate reads can be artifacts coming from the used experimental procedure (see Methods). Thus, to experimentally validate these insertion sites as hotspots, the six strongest resulting from the IntI1 experiment (corresponding to insertions in *ybhO*, *alsB*, *ilvD*, *pyrE*, *metC* and *yjhH* genes) were cloned in a recipient plasmid (Figure 5A). As a control, we used a genomic site used only once by IntI1 for cassette insertion, called US-*ygcE* (US for Unique Spot) and another site used 15 times, *i.e*., very close to the median of insertion site usage, called MIS-*abgA* (MS for Median Spot). In each case, the cloned segment encompassed the 300 and 200 bps flanking the 5’GWT3’ insertion point. The six hotspots showed a high rate of recombination (around 10^-3^, close to the classical *attI* and *attC* sites), while no recombination event was detected for the US and MS control sites (Figure 5A). Notably, the highest insertion efficiency was obtained for the *ybhO* hotspot, the one showing the highest insertion usage (209,387). These results confirm that these hotspots are regions attracting cassette insertions and not the consequences of experimental bias.

**Figure 5:**
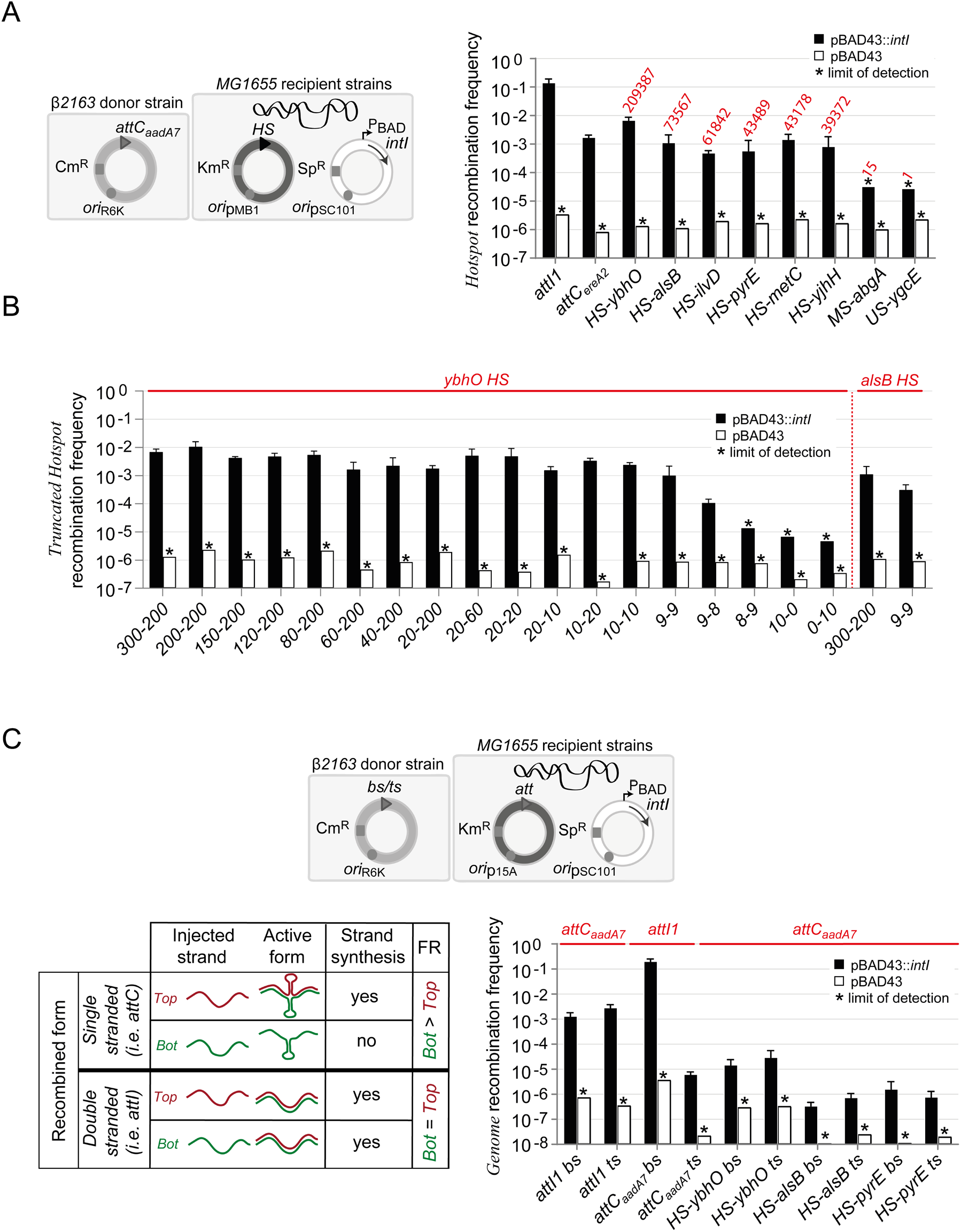
*Hotspot* sites as donor and receptor sites during conjugation assay. A) Testing the *hotspots* as receptor sites The experimental setup is described (left panel). The setup is the same as in Figure 3A, except that the donor plasmid contains an *attC_aadA7_* site and the MG1655 *E. coli* recipient strains contain recipient plasmids carrying *attI1, attC_ereA2_*, hotspot (HS), median spot (MS) and unique spot (US) sites (black triangle) and a high copy number origin of replication (*ori*pMB1). All recombinants are selected on Cm containing plates. The graph representing all the recombination frequencies obtained into the *attI1* and *attC_ereA2_* sites, into *ybhO, alsB, ilvD, pyrE, metC, yjhH* hotspot sites (HS), into the *abgA* median site (MS) and into the *ygcE* unique site (US) is shown (Right panel). The receptor plasmids are indicated in the axis-x legends (see also Table S1). The number of insertions that we previously obtained in each hotspot site (Figure 4A) are indicated in red at the top of the bars. B) Testing the truncated *hotspots* as receptor sites. The receptor plasmids are indicated in the axis-x legends (see table S2). The 2 numbers represent the number of base pairs kept on each 5’ and 3’ side of the cutting position in the *ybhO* and *alsB* hotspot sites (see Figure 4D). For more details on the calculation of the recombination frequency see the legend of the figure 3B. C) Testing the *hotspots* as donor sites. The experimental setup is described (left panel). The setup is the same as in Figure 3A, except that the donor plasmids carry different *att* and *hotspot* sites (grey triangle) delivering either the bottom (bs) or the top strand (ts) depending on the orientation of the recombination sites. The MG1655 *E. coli* recipient strains contain recipient plasmids carrying *attI1* or *attC_ereA2_* sites. All recombinants are selected on Cm containing plates. The graph representing all the recombination frequencies obtained into the *attI1* and *attC_ereA2_* sites, injecting top or bottom strands of the several donor sites is shown (Right panel). The donor plasmids are indicated in the axis-x legends and the receptor sites in red at the top of the bars (see also Table S1). Scheme of expected frequencies injecting the bottom or the top strands in function of the active recombination forms is shown. Bottom strand (bs, green line) or top strand (ts, red line) are injected. If the recombination occurs by a single-stranded bottom form, a strand synthesis (dotted line) is necessary only when the top strand is injected and not the bottom strand. The frequency of recombination (FR) is therefore expected higher injecting the bottom strand than the top one (Bot > Top). If recombination occurs by a double-stranded form, a strand synthesis (dotted line) is necessary injecting either of the two strands. The frequency of recombination is expected to be the same injecting the bottom strand or the top one (Bot = Top).

### The properties of genome insertion sites differ from *att* recombination sites

To further characterize the properties of genome sites, we used the *ybhO* hotspot site as proxy of insertion sites and determined its minimal functional length. We constructed and tested several lengths of base pairs on each 5’ and 3’ side of the cutting position of the *ybhO* hotspot site (Figure 5B). Decreasing the lengths to 9 nucleotides on each side maintained the recombination frequency above 10^-3^ (9-9, Figure 5B), defining a minimal 18 nt length-functional insertion, while smaller lengths hampered the recombination frequencies (Figure 5B). These functional 9-9 lengths were confirmed for the *alsB* hotspot (Figure 5B). Alignment of the 10 base pairs on each 5’ and 3’ side of the cutting position of the six hotspots from Figure 5A uncovered a consensus sequence, keeping the highly conserved 5’GWT3’ residues in positions -1 to 2, but revealing two supplementary motifs, on either side of the cleavage site (Figure S8A). We validated the importance of the 5’CRGM3’ right motif in positions 6 to 9. Indeed, replacing this motif by the 5’CCAG3’ and 5’TCAG3’ motifs in the *ybhO* hotspot decreased insertion frequencies by more than 2 logs compared to the *wt ybhO* 5’CAGC3’ or the consensus 5’CAGA3’ sequences (Figure S8B). Interestingly, performing the same position replacements in the right part of the *attI1* site did not hamper the recombination frequency (Figure S8C), indicating structural differences between the *attI* and genome sites. From these, we conclude that genome sites differ broadly, in terms of sequence and structure from both classical *attC* and *attI* recombination sites. We named these new integron sites *“attG”*, where G stands for “genome”.

### *attG* sites are recombined as double-stranded forms and are single-cleaved by the integrase

To determine the double-stranded or the single-stranded nature of the *attG* sites, we cloned three hotspots in the donor plasmid, in both orientations, delivering either the top or the bottom strand as donor sites during conjugation, and we tested the ability of these *attG* sites to recombine in an *attC_aadA7_* receptor site cloned in the recipient plasmid (Figure 5C). If *attG* sites are recombined as a strand specific single-stranded form, a difference in the recombination rate would be expected while injecting the top or the bottom strand as shown for *attC* sites ((8), Figure 5C). In contrast, if *attG* are recombined as a double-stranded form and thus requires second strand synthesis to be effective, no difference of recombination frequency is expected whatever the injected strand as shown for the *attI* site ((8), Figure 5C). Obtained insertion frequencies varied between the tested hotspots but were similar regardless of their orientation (Figure 5C). These results confirm that *attG* sites are recombined as double-stranded form, like the *attI* sites. PCR and sequencing of recombined products demonstrated a sequence homogeneity around the insertion site thus confirming that recombination occurs by a single cleavage of the doubled-stranded matrix (Figure S9A). Indeed, a double cleavage would lead to a heterogeneity of sequences at the insertion point as expected when the core sequences of *att* sites are different (Figure S19A). We also confirmed by sequence analysis and determination of the cassette insertion orientation that recombination takes place between the bottom strands of both the hotspots and *attC* receptor sites and at the expected recombination points (Figure S9B). Note that we sequenced more than 24 recombination products for each top and bottom injected strands and for all three tested hotspot sites (more than 144 sequences total), illustrating the robustness of our results. We therefore conclude that during cassette insertion in the genome, the *attG* sites recombine as double-stranded forms and that only their bottom strands are cleaved.

## DISCUSSION

Most studies on integrons focused on the recombination properties of cassettes in the proper integron *att* sites. However, a few experiments revealed the ability of cassettes to insert at a very low rate (around 10^-6^) in secondary sites in plasmids and genomes (23–27). Moreover, the single and complete *aadB* cassette has been found inserted at a secondary site into the IncQ plasmid RSF1010 (28) and into the pRAY plasmid (29,30) just downstream of a known promoter ensuring the expression of the *aadB* antibiotic resistance gene. These rare insertion events have been overlooked for decades and considered anecdotal in the integron functioning. Here, by performing an extensive bioinformatics analysis of available sequenced bacterial genomes, we found many isolated cassettes inserted in bacterial genomes. We called these cassettes, SALIN, for Single *attC* sites lacking integron-integrase and we took this observation as a starting point to study and identify a so far hidden cassette dissemination route in bacterial genomes. Using an assay mimicking the natural conditions in which the acquisition of cassettes occurs through horizontal gene transfer, we demonstrated that integron cassettes can disseminate outside the integron platform, at several positions in host genomes and at a high frequency, *i.e*., close to the frequency obtained using canonical *attI and attC* sites. We unveiled that integrase insertion sites have a very small 5’GWT3’ consensus sequence meaning that the cassette insertional landscape can be very large. As example, in the 4,641,652 bps of the MG1655 *E. coli* genome, this represents 338,348 theoretically targetable sites. We demonstrated that cassette insertions can induce genome changes by disrupting genes and that excision of genome inserted cassettes, when occurring outside the insertion site, can result in the deletion of pieces of the genome. Cassette insertion can also represent a gain of function for the bacteria when genome inserted cassettes are expressed. These cassettes could also be domesticated since the folded single-stranded *attC* sites can be easily deleted by replication slippage events (31,32).

Interestingly, another site-specific recombinase, the lambda (λ) integrase, which ensure λ phage genome integration at the *attB* specific site in the host chromosome, was also described as being able to catalyze phage genome integration into genome sites. However, the observed consensus insertion sequence is much larger than that of integron integrases restraining the secondary insertion sites to a small number of locations in bacterial chromosome (33,34). Therefore, compared to the integron integrase, genome insertions mediated by the λ integrase would impact very poorly the genome. Another difference is that the secondary sites of the λ integrase are actually very similar to the *attB* recombination site, probably resulting from an off-target activity of the λ integrase. On the contrary, we have shown that the genomic sites of the integrase are structurally different from both *attC* and *attI* sites, i.e., with a cleavage point located in their central part, thus bringing them closer to classical double-stranded core recombination sites such as *dif* sites (35). We therefore called these new sites, *attG* sites. As these *attG* sites are recombined as a double-stranded form, we suggest that *attI* sites could have derived from the sequence of one of these hotspots through coevolution with an integrase.

In conclusion, we have demonstrated the existence of a new cassette recombination route that greatly expands the role of integrons in the dissemination of adaptive functions such as antibiotic resistance (Figure 6). This new route could allow bacteria to “safeguard”cassettes in their genomes. This could be particularly advantageous in conditions where MIs are carried by conjugative plasmids that cannot be maintained in the cell. Beyond that, these results show that the integron system could represent a general mechanism for genomic diversification driving bacterial evolution.

**Figure 6:**
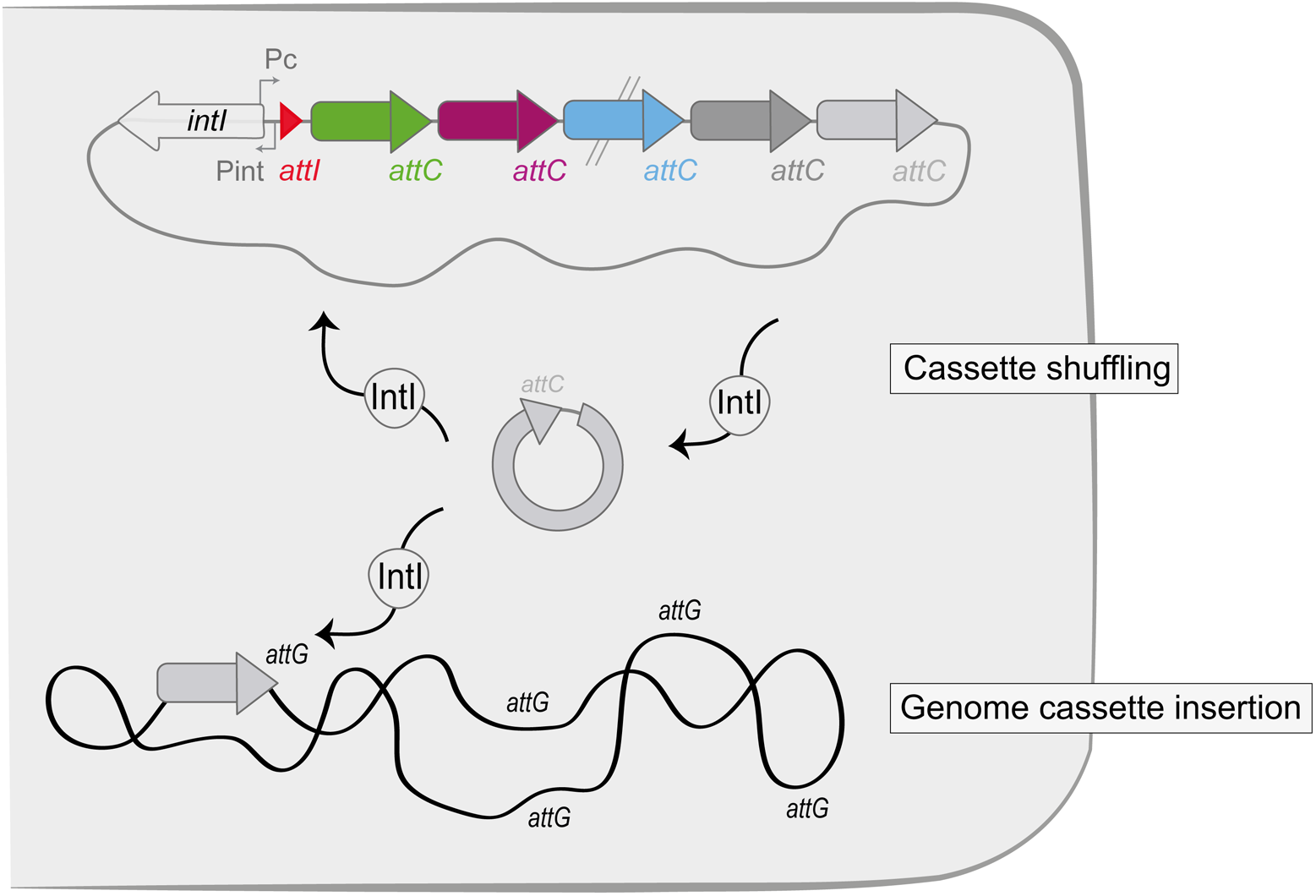
The new route of the integron cassette dissemination. Upon expression of the integrase, cassette shuffling inside the integron can occur through cassette excision (*attC* × *attC*) and insertion of the excised cassettes in the first position in the array (*attI* × *attC*). Cassettes can also be inserted in bacterial genomes through *attC* × *attG* recombination.

## SUPPLEMENTAL INFORMATION

Table S1: Bacterial strains, plasmids and primers used in this study

Figure S1: Random PCR approach used to determine the genome insertion sites

Figure S2: Cassette insertion events in genome in presence of an integron

Figure S3: Cassette recombination events mediated by IntI2 and IntI3 integrases

Figure S4: Representation of the imprecise excisions of genome-inserted cassettes

Figure S5: Description of the library construction for Deep sequencing

Figure S6: Computational analysis of Deep sequencing data

Figure S7: Digital PCR analysis of the *oriC* copy number relative to *terC* in *E. coli* receptor strain during a conjugation mimicking assay

Figure S8: Analysis of the *attG* insertion site motifs

Figure S9: Determination of the *attG* recombination nature

## ACKNOWLEDGEMENTS

We would like to thank Gaspard Macaux for its experimental help. We also thank all the lab members for helpful discussion.

## AUTHOR CONTRIBUTIONS

C.L and DM designed the research. C.L, E.R, C.V, B.D, D. L, V.P, F.L and T.N performed the experiments. G.M and F. L performed the computational analysis of Deep sequencing data. E.L, J. C and E.P.C.R performed the bioinformatics genomics analysis. C.L and G. M wrote the draft of the manuscript. All authors read, amended the manuscript, and approved its final version.

