## Supplemental figures and table for "A new route for integron cassette dissemination among bacterial genomes"

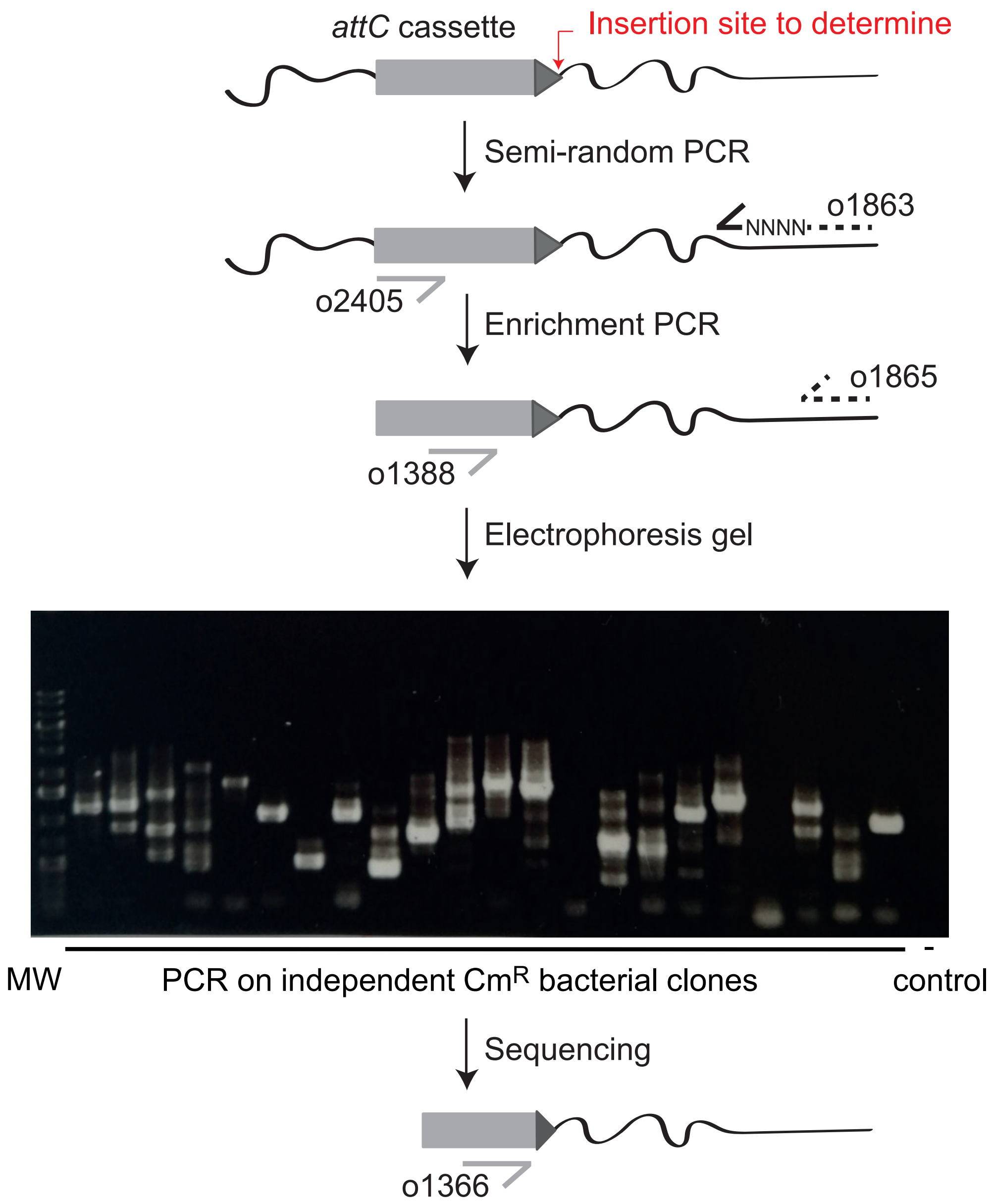


**Figure S1: Random PCR approach used to determine the genome insertion sites**
The genome inserted *attC* cassette is represented by a light grey arrow (CDS) followed by a dark grey triangle (*attC*). The red arrow indicates the insertion site to determine. Primers used for the PCR ampification and sequencing are shown. Electrophoresis analysis of PCR products is shown. MW: Molecular Weight Marker

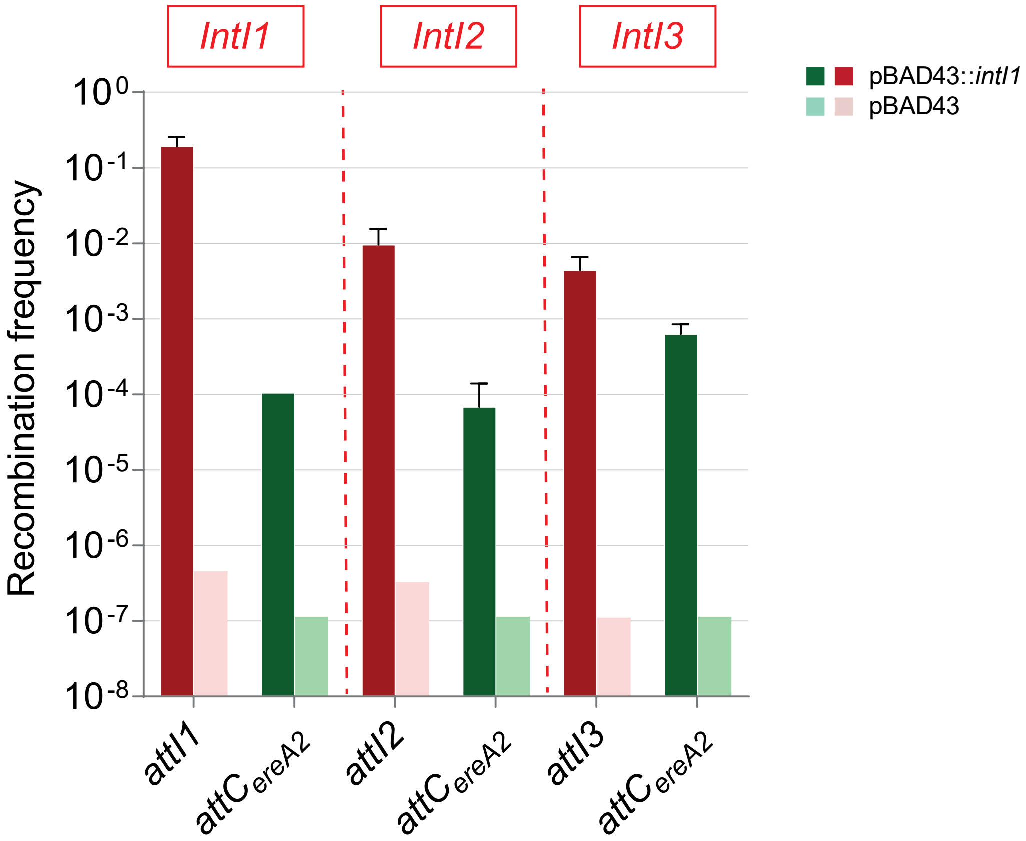


**Figure S2: Cassette recombination events mediated by IntI1, IntI2 and IntI3 integrases**
Frequencies of *aadA7* cassette insertion into the *attI* sites (red and pink bars) and *attC*$_{ereA2}$ (light and dark green bars) sites are represented.

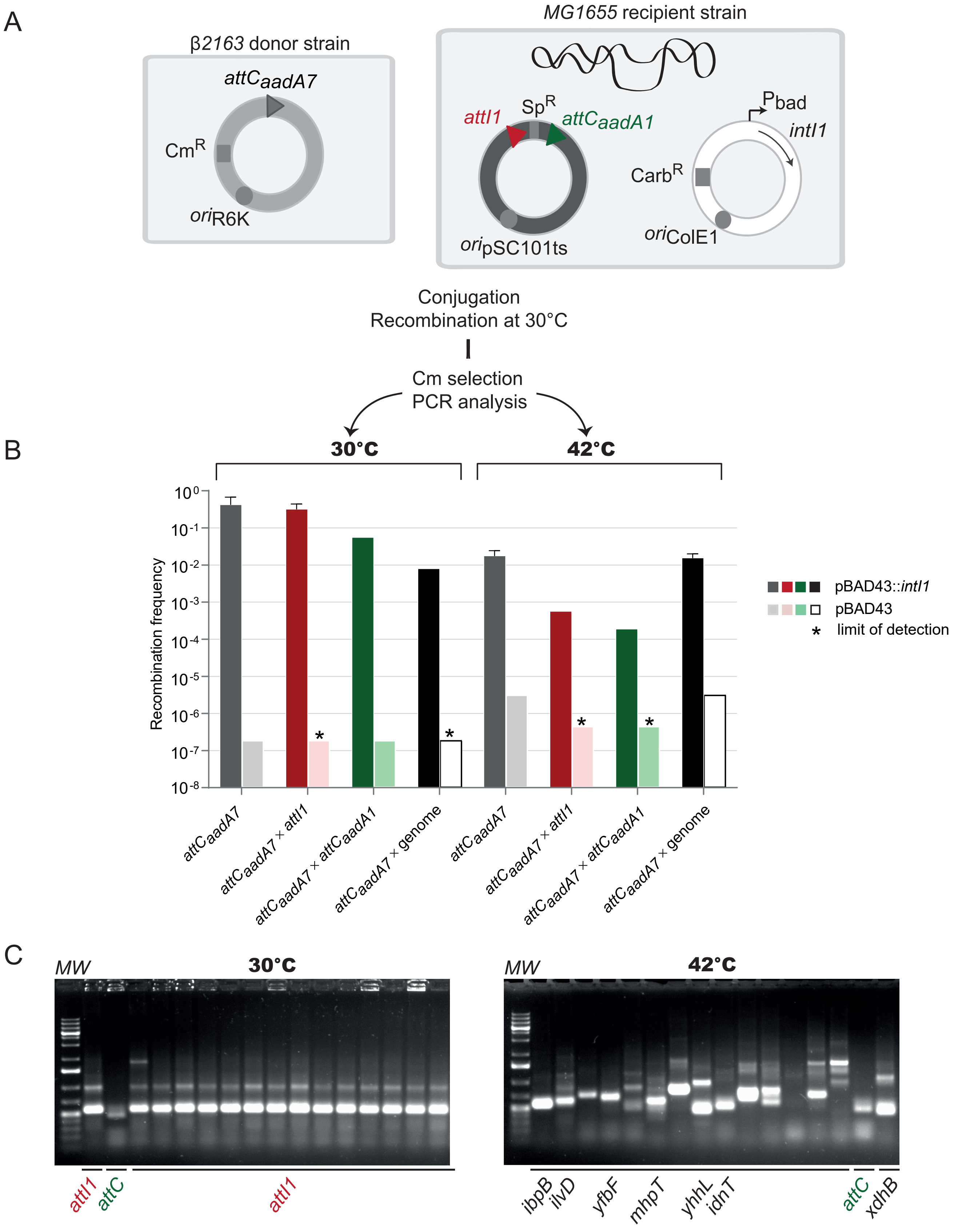


**Figure S3: Cassette insertion events in genome in presence of an integron**

(A) Experimental setup of the cassette insertion assay

See legend of the Figure 3A for details. ts: thermosensitive

(B) Frequency of cassette insertion into the *att* sites and genome

(C) Electrophoresis migration of PCR products

Several PCR products are sequenced and the insertion sites are indicated (*att* sites and gene location). MW: Molecular Weight Marker

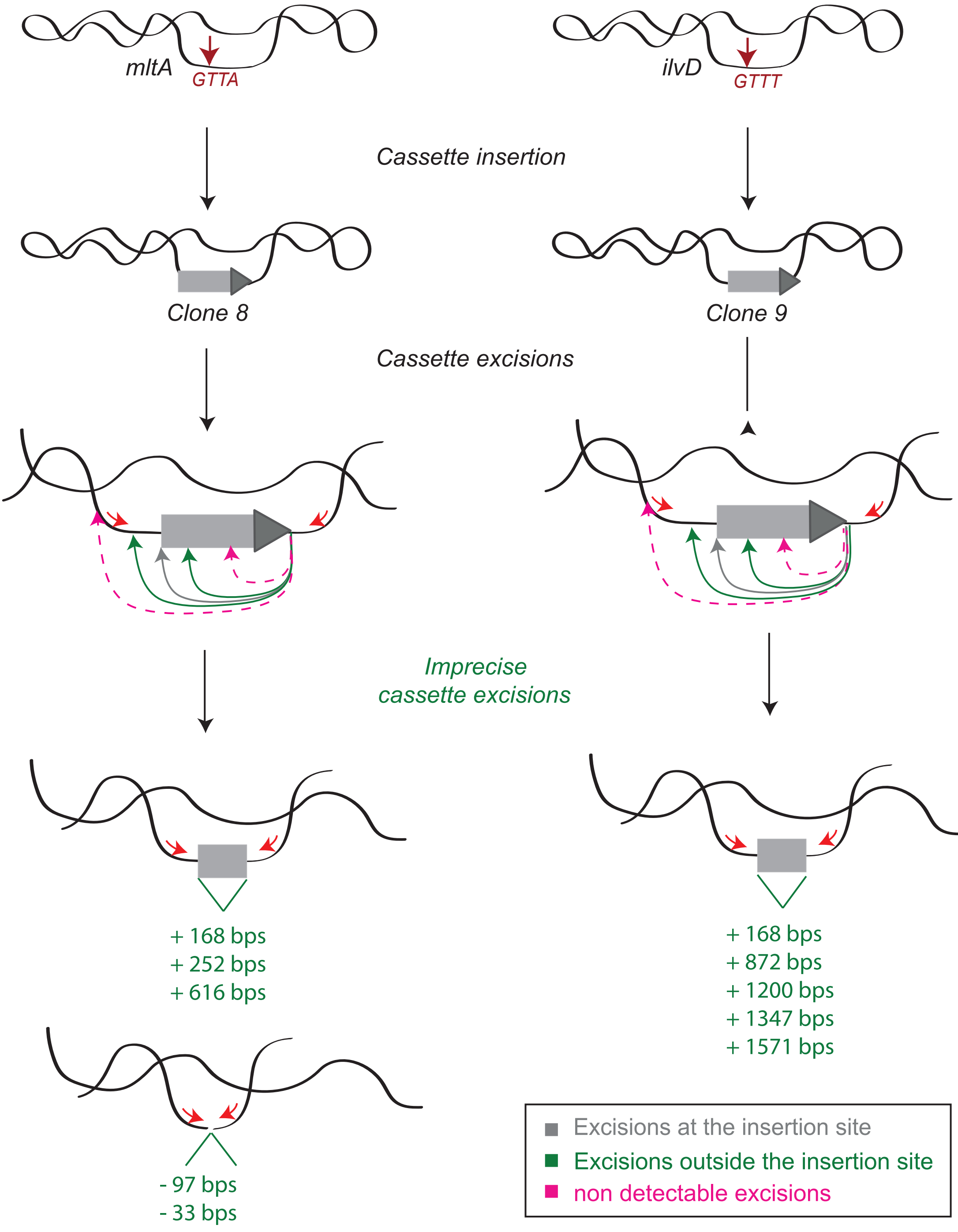


**Figure S4: Representation of the excisions of genome inserted cassettes**
Primers used to perform PCR detecting the excision events are shown by red arrows. Non detectable imprecise excisions correspond to excision that can not be revealed by PCR using our set of primers. +, means that cassette excision point is localized inside the inserted cassettes (adding DNA sequence) and -, outside (removing bacterial genome DNA).

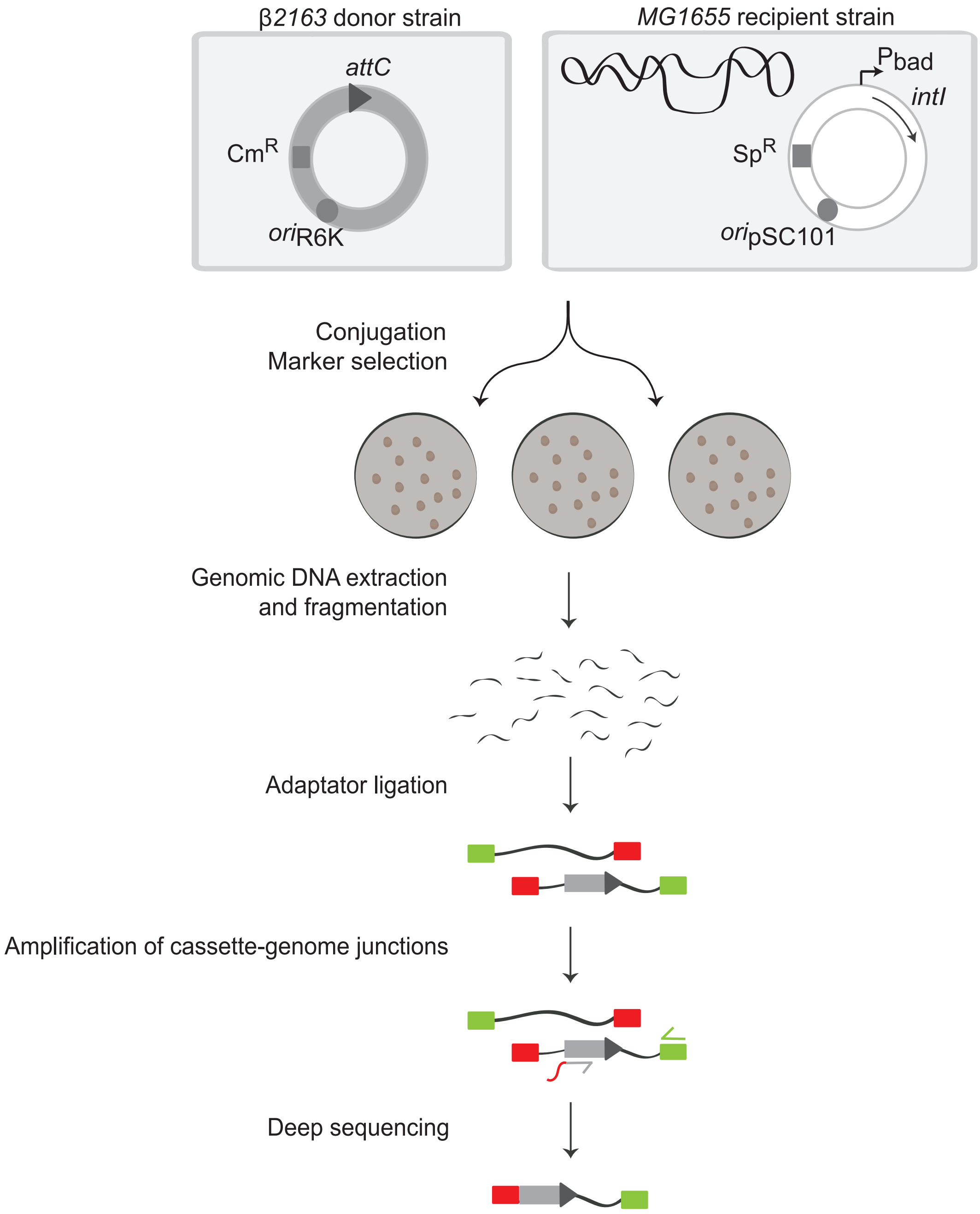


**Figure S5: Description of the library construction for Deep sequencing**
All the steps of the library construction are indicated. Adaptators are represented by red and green rectangles. The *attC* cassette is represented by a light grey arrow (CDS) followed by a dark grey triangle (*attC*).

| Int | Collected clones | Total reads | Reads after duplicate removal | Insertion sites |
|---|---|---|---|---|
| IntI1 | 49,600 | 3,205,043 | 361,464 | 22,271 |
| IntI2 | 117,048 | 6,813,145 | 154,246 | 9,125 |
| IntI3 | 349,920 | 6,584,879 | 210,104 | 19, 287 |

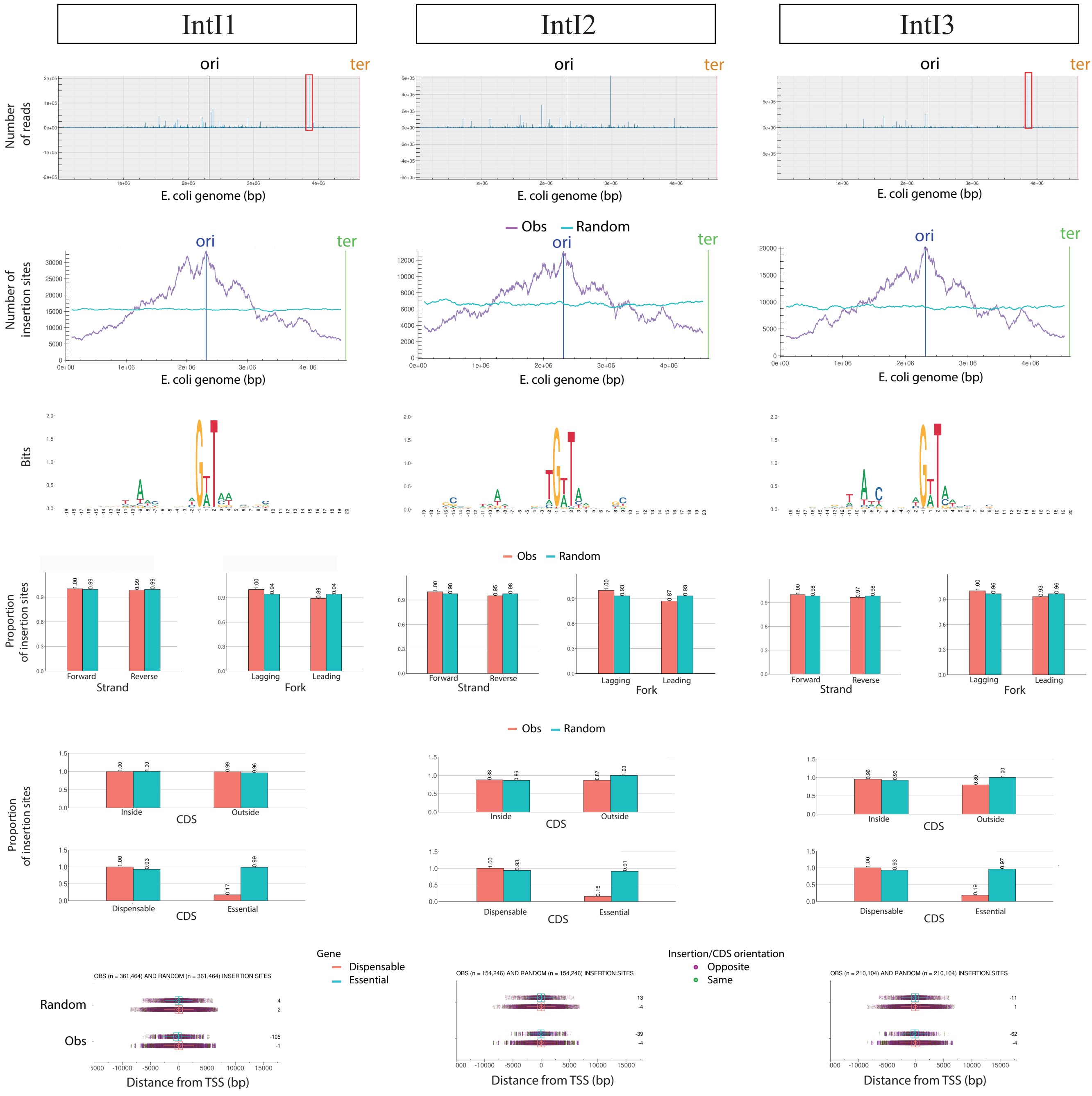


**Figure S6: Computational analysis of Deep sequencing data**

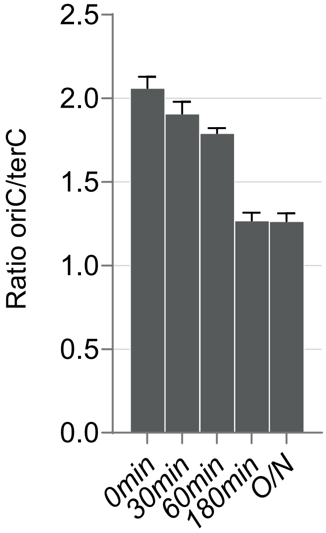


**Figure S7: Digital PCR analysis of the oriC copy number relative to terC (Ratio oriC/terC) in *E. coli* receptor strain during a conjugation mimicking assay** DNA extraction and PCR were performed at 0, 30, 60, 180 minutes (min) after the beginning of the conjugation and also after an overnight (O/N) time of incubation. Note that only receptor strains are used during this assay mimicking conjugation.

A

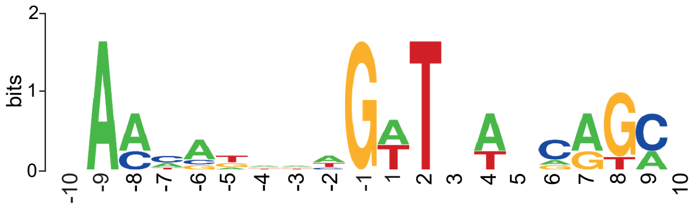


B

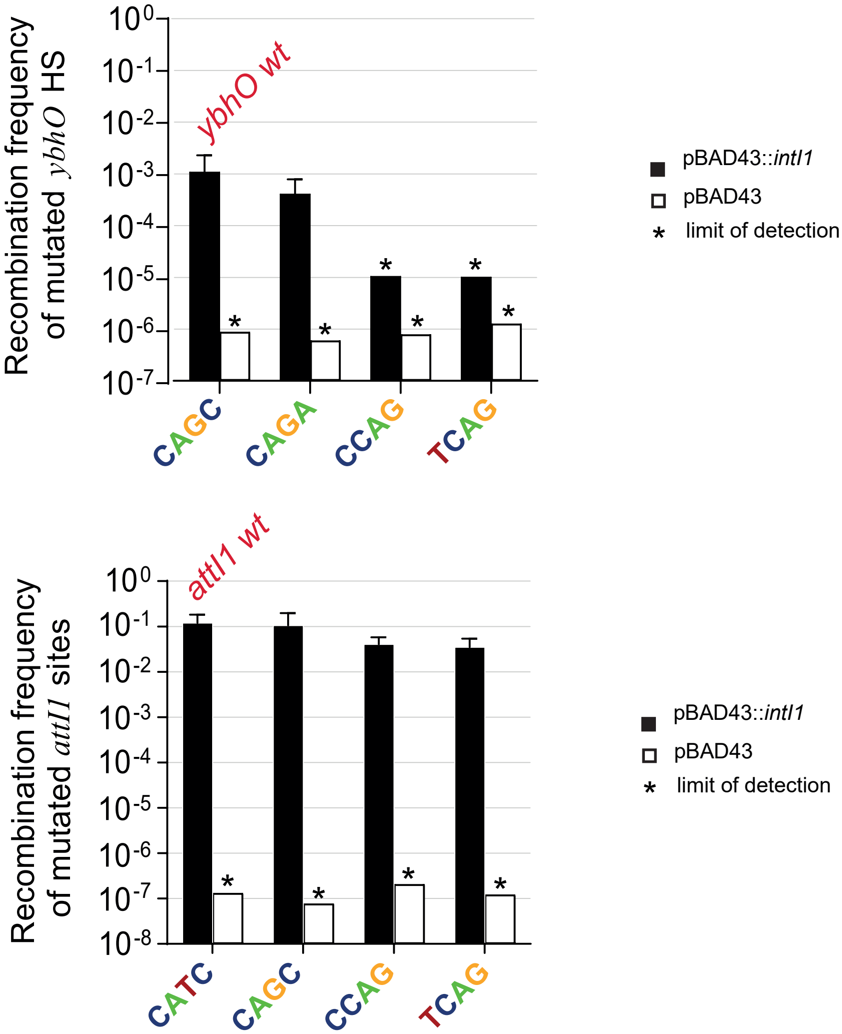


**Figure S8: Analysis of the *attG* insertion site motifs**
(A) Consensus sequence of the six highest hotspot insertion sites. A total of 20 bases around the cleavage point was inputted into the WebLogo program (http://weblogo.berkeley.edu/) to generate the motif. The cleavage occurs between the -1 and 1 bases.
Bits refers to the information content.
(B) Frequencies of cassette insertion into the mutated *ybhO* hotspot (HS) and *attI1* sites.
The indicated 4 nt sequences correspond to the position 6 to 9 given that position -1 corresponds to the G of the cleaved triplet site (see also S9A).

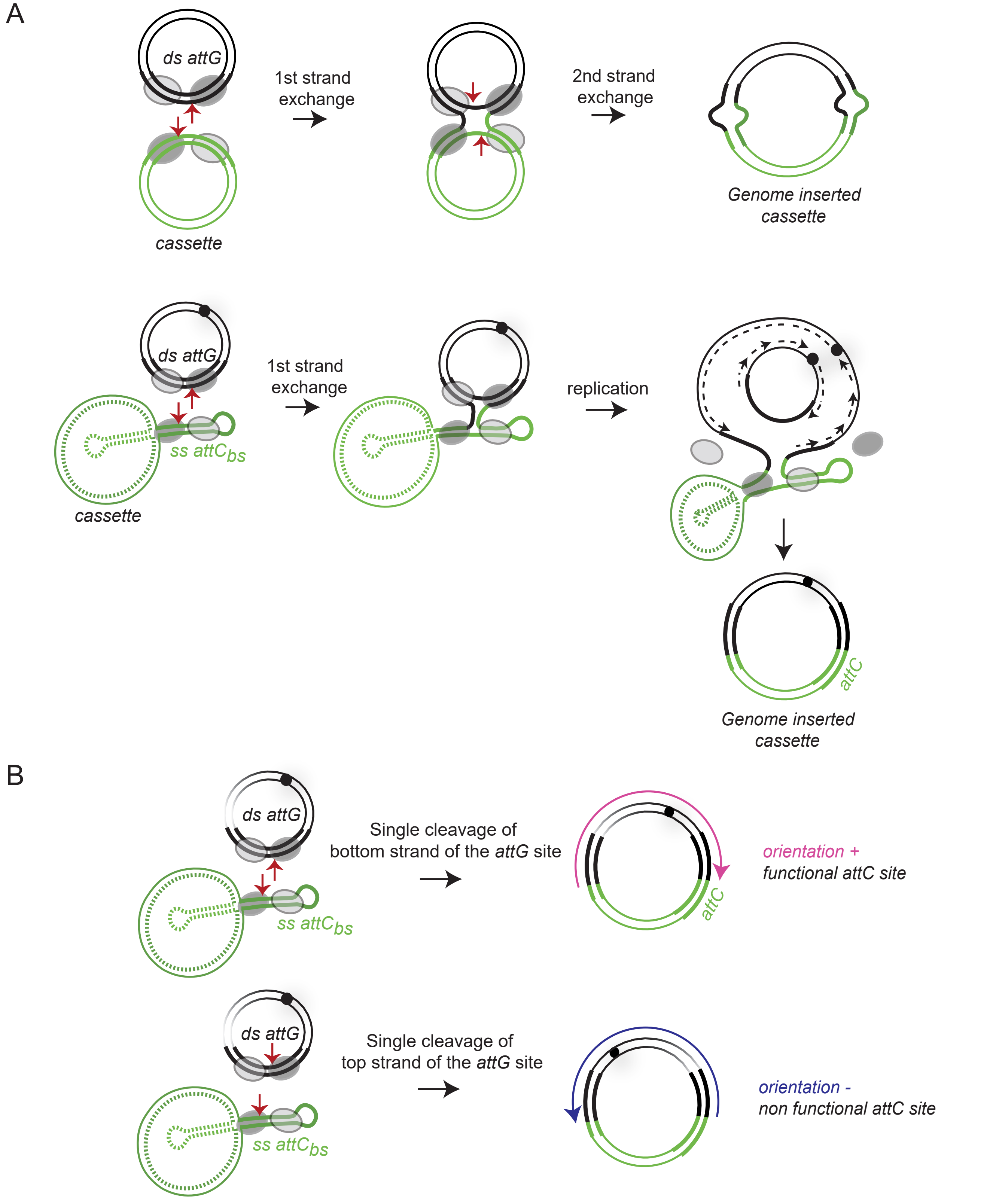


**Figure S9: Determination of the *attG* recombination nature**

(A) Double cleavage of *attG* sites: we expect heterogeneity of products due to the repair process.
Single cleavage of *attG* sites: we expect homogeneity of products due to the replication.

(B) Single cleavage of bottom or top strands (bs and ts) of *attG* sites

The (+) orientation (pink arrow) results from a bottom strand cleavage while the (-) (blue arrow) results from a top strand cleavage.

*attG* containing replicons are represented by dark lines and *attC* cassettes by green ones.

The precise cleavage point is indicated by a red arrowhead.

### Table S1: Bacterial strains, plasmids and primers used in this study

### Bacterial strains

| Strain number | Relevant genotypes or description | References |
|---|---|---|
| | ***Bacterial strains*** | |
| 8195 | DH5α (F-) *supE44 ΔlacU169* (ϕ80lacZΔM15) *ΔargF hsdR17 recA1 endA1 gyrA96 thi-1 relA1* | Laboratory collection |
| One Shot™TOP10 | Top10 (F-) *mcrA* Δ( *mrr-hsd*RMS-*mcr*BC) Φ80*lac*ZΔM15 Δ *lac*X74 *rec*A1 *ara*D139 Δ( *araleu*) 7697 *gal*U *gal*K *rps*L *end*A1 *nup*G [$Str^R$] | Thermo Fischer Scientific (Ref: C404010) |
| 9420 | π1, DH5α *ΔthyA*::(*erm-pir116*) [$Erm^R$] | Demarre, et al. 2005 |
| 8725 | π3813 *lacIq thi-1 supE44 endA1 recA1 hsdR17 gyrA462 zei-298::Tn10 ΔthyA*::(*erm-pir116*) [$Erm^R$] | Le Roux, et al. 2007 |
| 4196 | β2163 (F-) RP4-2-Tc::Mu *ΔdapA*::(*erm-pir*) [$Km^R$ $Erm^R$] | Demarre, et al. 2005 |
| 8726 | β3914, β2163 *gyrA462 zei-298::Tn10* [$Km^R$ $Erm^R$] | Le Roux, et al. 2007 |
| C349 | MG1655 *E. coli* K12 *wt* | Laboratory collection |
| 4826 | MG1655 *recA269::Tn10* [$Tc^R$] | RG Lloyd |
| K779-K781 | N16961 *V. cholerae ΔattIA* | Vit, et al. 2021 |
| | **Escherichia coli *transformed donor strains for suicide conjugation assay*** | |
| D059 | β2163 pD059 | Nivina, et al. 2016 |
| L120 | β2163 pD060 | Nivina, et al. 2016 |
| F487 | β2163 pF486 | This study |
| 3651 | β2163 p3614 | This study |
| 3653 | β2163 p3615 | This study |
| 3652 | β2163 p3616 | This study |
| 2714 | β2163 p2714 | Bouvier, et al. 2005 |
| 2715 | β2163 p2715 | Bouvier, et al. 2005 |
| Q997-Q998 | β2163 pO323-pO324 | This study |
| O344-O346 | β2163 pO321-pO322 | This study |
| O788-O789 | β2163 pO749-pO750 | This study |
| O790-O791 | β2163 pO751 | This study |
| O792-O793 | β2163 pO752 | This study |
| O794-O795 | β2163 pO753-pO755 | This study |
| N147-N698-N699 | β2163 pN695-pN697 | This study |
| N876-N881 | β2163 pN705-pN707 | This study |
| N882-N885 | β2163 pN708-pN709 | This study |
| M787-M789 | β3914 pM779-pM781 | This study |
| | **Escherichia coli *transformed receptor strains for suicide conjugation assay*** | |
| L503-L504 | MG1655 pL290 | This study |
| L501-L502 | MG1655 pL294 | This study |
| O859-O860 | MG1655 pO859-pO860 | This study |
| O727-O728 | MG1655 pO727-pO728 | This study |
| L385-L387 | MG1655 p929 pL290 | This study |
| J733-J735 | MG1655 p929 pL294 | This study |

| | | |
|---|---|---|
| O889-O891 | MG1655 pO846 pL290 | This study |
| O880-O882 | MG1655 pO846 pO859 | This study |
| O892-O894 | MG1655 pO847 pL290 | This study |
| O886-O888 | MG1655 pO847 pO727 | This study |
| L515-L516 | MG1655 pG164 pL290 | This study |
| L512-L513 | MG1655 pG164 pL294 | This study |
| O877-O879 | MG1655 pG164 pO859 | This study |
| O883-O885 | MG1655 pG164 pO727 | This study |
| M371-M373 | MG1655 pM335 p979 | This study |
| M368-M370 | MG1655 pM335 p3938 | This study |
| N463-N465 | MG1655 *recA* pN435-pN437 | This study |
| N460-N462 | MG1655 *recA* pN438-pN440 | This study |
| T125-T127 | MG1655 pS957 pL290 | This study |
| T128-T130 | MG1655 pS957 pL294 | This study |
| T428-T430 | MG1655 pT370 pL290 | This study |
| T431-T433 | MG1655 pT370 pL294 | This study |
| O156-O157 | MG1655 pO122 pL290 | This study |
| O176-O177 | MG1655 pO122 pL294 | This study |
| O158-O159 | MG1655 pO123 pL290 | This study |
| O178-O179 | MG1655 pO123 pL294 | This study |
| O318-O320 | MG1655 pO343 pL290 | This study |
| O315-O317 | MG1655 pO343 pL294 | This study |
| O162-O163 | MG1655 pO125 pL290 | This study |
| O182-O183 | MG1655 pO125 pL294 | This study |
| O164-O165 | MG1655 pO126 pL290 | This study |
| O184-O185 | MG1655 pO126 pL294 | This study |
| O166-O167 | MG1655 pO127 pL290 | This study |
| O186-O187 | MG1655 pO127 pL294 | This study |
| O395-O396 | MG1655 pO382 pL290 | This study |
| O403-O404 | MG1655 pO382 pL294 | This study |
| O401-O402 | MG1655 pO385 pL290 | This study |
| O409-O410 | MG1655 pO385 pL294 | This study |
| O772-O773 | MG1655 pO721 pL290 | This study |
| O774-O775 | MG1655 pO721 pL294 | This study |
| P116-P118 | MG1655 pO743 pL290 | This study |
| P119-P121 | MG1655 pO743 pL294 | This study |
| O776-O777 | MG1655 pO723 pL290 | This study |
| O778-O779 | MG1655 pO723 pL294 | This study |
| O780-O781 | MG1655 pO725 pL290 | This study |
| O782-O783 | MG1655 pO725 pL294 | This study |
| O996-O998 | MG1655 pO970 pL290 | This study |
| O999-P002 | MG1655 pO970 pL294 | This study |
| P003-P005 | MG1655 pO971 pL290 | This study |
| P006-P008 | MG1655 pO971 pL294 | This study |
| P122-P124 | MG1655 pO974 pL290 | This study |
| P125-P127 | MG1655 pO974 pL294 | This study |
| P282-P284 | MG1655 pP177 pL290 | This study |
| P285-P286 | MG1655 pP177 pL294 | This study |
| P288-P290 | MG1655 pP178 pL290 | This study |
| P291-P293 | MG1655 pP178 pL294 | This study |

| | | |
|---|---|---|
| P916-P918 | MG1655 pP761 pL290 | This study |
| P919-P921 | MG1655 pP761 pL294 | This study |
| P928-P930 | MG1655 pP807 pL290 | This study |
| P931-P933 | MG1655 pP807 pL294 | This study |
| P922-P924 | MG1655 pP762 pL290 | This study |
| P925-P927 | MG1655 pP762 pL294 | This study |
| Q363-Q365 | MG1655 pQ223 pL290 | This study |
| Q366-Q368 | MG1655 pQ223 pL294 | This study |
| Q375-Q377 | MG1655 pQ342 pL290 | This study |
| Q378-Q380 | MG1655 pQ342 pL294 | This study |
| Q369-Q371 | MG1655 pQ162 pL290 | This study |
| Q372-Q374 | MG1655 pQ162 pL294 | This study |
| Q381-Q383 | MG1655 pQ159 pL290 | This study |
| Q384-Q386 | MG1655 pQ159 pL294 | This study |
| P910-P912 | MG1655 pP760 pL290 | This study |
| P913-P915 | MG1655 pP760 pL294 | This study |
| Q728-Q730 | MG1655 pQ706 pL290 | This study |
| Q731-Q733 | MG1655 pQ706 pL294 | This study |
| R149-R151 | MG1655 pR102 pL290 | This study |
| R152-R154 | MG1655 pR102 pL294 | This study |
| R155-R157 | MG1655 pR105 pL290 | This study |
| R158-R160 | MG1655 pR105 pL294 | This study |
| R069-R071 | MG1655 pQ980 pL290 | This study |
| R072-R074 | MG1655 pQ980 pL294 | This study |
| R081-R083 | MG1655 pR010 pL290 | This study |
| R084-R086 | MG1655 pR010 pL294 | This study |
| R087-R089 | MG1655 pR012 pL290 | This study |
| R090-R092 | MG1655 pR012 pL294 | This study |
| R093-R095 | MG1655 pR062 pL290 | This study |
| R096-R098 | MG1655 pR062 pL294 | This study |
| R568-R569 | MG1655 pR528 pL290 | This study |
| R570-R571 | MG1655 pR528 pL294 | This study |
| O373-O375 | MG1655 *recA* pO371-pO372 pL290 | This study |
| O377-O379 | MG1655 *recA* pO371-pO372 pL294 | This study |
| O635 | MG1655 *recA inserted cassette clone 7* pN435 | This study |
| O636 | MG1655 *recA inserted cassette clone 8* pN435 | This study |
| O637 | MG1655 *recA inserted cassette clone 9* pN435 | This study |
| O638 | MG1655 *recA inserted cassette clone 7* pM889 | This study |
| O639 | MG1655 *recA inserted cassette clone 8* pM889 | This study |
| O640 | MG1655 *recA inserted cassette clone 9* pM889 | This study |
| O641 | MG1655 *recA inserted cassette clone 7* pM888 | This study |
| O642 | MG1655 *recA inserted cassette clone 8* pM888 | This study |
| O643 | MG1655 *recA inserted cassette clone 9* pM888 | This study |
| **Vibrio cholerae *transformed receptor strains for suicide conjugation assay*** | | |
| L397-L399 | N16961 *ΔattIA* pL290 | This study |
| L400-L402 | N16961 *ΔattIA* pL294 | This study |

| **Plasmid number** | **Plasmid description** | **Relevant properties and construction** |
|---|---|---|

| | | |
|---|---|---|
| p929 | pSU38Δ::*attI1* | *ori*p15A; [$Km^R$] (Biskri et al. 2005) |
| pQ980 | pSU38Δ::*attI1-BamHI* | *ori*p15A; [$Km^R$], amplification by inverse PCR of p929 with primers o6593 and o6594. |
| pR010 | pSU38Δ::*attI1-CCAG* | *ori*p15A; [$Km^R$], amplification by Quick change PCR of pQ980 with primers o6605 and o6606. |
| pR012 | pSU38Δ::*attI1-TCAG* | *ori*p15A; [$Km^R$], amplification by Quick change PCR of pQ980 with primers o6607 and o6608. |
| pR062 | pSU38Δ::*attI1-CAGC* | *ori*p15A; [$Km^R$], amplification by inverse PCR of pQ980 with primers o6919 and o6920. |
| pO846-pO861 | pSU38Δ::*attI2* | *ori*p15A; [$Km^R$], annealing of complementary o6314 and o6315 primers reconstituting the *attI2* site. The annealed primers (generating digested *EcoRI* and *BamHI* enzyme restriction sites) are cloned in *EcoRI*/*BamHI* digested p929. |
| pO847-pO848 | pSU38Δ::*attI3* | *ori*p15A; [$Km^R$], annealing of complementary o6316 and o6317 primers reconstituting the *attI3* site. The annealed primers (generating digested *EcoRI* and *BamHI* enzyme restriction sites) are cloned in *EcoRI*/*BamHI* digested p929. |
| pG164 | pSU38Δ::$attC_{ereA2}$ | *ori*p15A; [$Km^R$], annealing of complementary o4870 and o4871 primers reconstituting the $attC_{ereA2}$ site. The annealed primers (generating digested *EcoRI* and *BamHI* enzyme restriction sites) are cloned in *EcoRI*/*BamHI* digested p929. |
| pO371-pO372 | pSU38Δ::$attC_{aadA7}$ | *ori*p15A; [$Km^R$], annealing of complementary o3849 and o3850 primers reconstituting the $attC_{aadA7}$ site. The annealed primers (generating digested *EcoRI* and *BamHI* enzyme restriction sites) are cloned in *EcoRI*/*BamHI* digested p929. |
| pS957-pS959 | pTOPO-*attI1* | *ori*MB1; [$Km^R$], the *attI1* fragment was amplified by PCR from pQ980 using o6699 and o6700 and cloned into the pCR®-BluntII-TOPO® vector. |
| pT370-pT371 | pTOPO-$attC_{ereA2}$ | *ori*MB1; [$Km^R$], annealing of complementary o6716 and o6717 primers reconstituting the $attC_{ereA2}$ fragment. The annealed primers (generating digested *EcoRI* enzyme restriction site) are cloned in *EcoRI* digested pCR®-BluntII-TOPO® vector. |
| pO122 | pTOPO-*HS-yjhH* | *ori*MB1; [$Km^R$], the HS *yjhH* fragment was amplified by PCR from MG1655 using o6273 and o6274 primers and cloned in the pCR®-BluntII-TOPO® vector. |
| pO123 | pTOPO-*HS-ybhO* | *ori*MB1; [$Km^R$], the HS *ybhO* fragment was amplified by PCR from MG1655 using o6275 and o6276 primers and cloned in the pCR®-BluntII-TOPO® vector. |
| pO343 | pTOPO-*HS-metC* | *ori*MB1; [$Km^R$], the HS *metC* fragment was amplified by PCR from MG1655 using o6277 and o6278 primers and cloned in the pCR®-BluntII-TOPO® vector. |
| pO125 | pTOPO-*HS-pyrE* | *ori*MB1; [$Km^R$], the HS *pyrE* fragment was amplified by PCR from MG1655 using o6279 and |

| | | |
|---|---|---|
| | | o6280 primers and cloned in the pCR®-BluntII-TOPO® vector. |
| pO126 | pTOPO-*HS-ilvD* | *ori*pMB1; [Km$^R$], the HS *ilvD* fragment was amplified by PCR from MG1655 using o6281 and o6282 primers and cloned in the pCR®-BluntII-TOPO® vector. |
| pO127 | pTOPO-*HS-alsB* | *ori*pMB1; [Km$^R$], the HS *alsB* fragment was amplified by PCR from MG1655 using o6283 and o6284 primers and cloned in the pCR®-BluntII-TOPO® vector. |
| pO382 | pTOPO-*MS-abgA* | *ori*pMB1; [Km$^R$], the MS *abgA* fragment was amplified by PCR from MG1655 using o6293 and o6294 primers and cloned in the pCR®-BluntII-TOPO® vector. |
| pO385 | pTOPO-*US-ygcE* | *ori*pMB1; [Km$^R$], the US *ygcE* fragment was amplified by PCR from MG1655 using o6299 and o6300 primers and cloned in the pCR®-BluntII-TOPO® vector. |
| pO721 | pTOPO-*HS-ybhO* (200(G)-200) | *ori*pMB1; [Km$^R$], the HS-*ybhO* (200(G)-200) fragment was amplified by PCR from MG1655 using o6276 and o6301 primers and cloned in the pCR®-BluntII-TOPO® vector. |
| pO743 | pTOPO-*HS-ybhO* (150(G)-200) | *ori*pMB1; [Km$^R$], the HS-*ybhO* (150(G)-200) fragment was amplified by PCR from MG1655 using o6276 and o6313 primers and cloned in the pCR®-BluntII-TOPO® vector. |
| pO723 | pTOPO-*HS-ybhO* (120(G)-200) | *ori*pMB1; [Km$^R$], the HS-*ybhO* (120(G)-200) fragment was amplified by PCR from MG1655 using o6276 and o6303 primers and cloned in the pCR®-BluntII-TOPO® vector. |
| pO725 | pTOPO-*HS-ybhO* (80(G)-200) | *ori*pMB1; [Km$^R$], the HS-*ybhO* (80(G)-200) fragment was amplified by PCR from MG1655 using o6276 and o6304 primers and cloned in the pCR®-BluntII-TOPO® vector. |
| pO970 | pTOPO-*HS-ybhO* (60(G)-200) | *ori*pMB1; [Km$^R$], the HS-*ybhO* (60(G)-200) fragment was amplified by PCR from MG1655 using o6276 and o6414 primers and cloned in the pCR®-BluntII-TOPO® vector. |
| pO971 | pTOPO-*HS-ybhO* (40(G)-200) | *ori*pMB1; [Km$^R$], the HS-*ybhO* (40(G)-200) fragment was amplified by PCR from MG1655 using o6276 and o6415 primers and cloned in the pCR®-BluntII-TOPO® vector. |
| pO974 | pTOPO-*HS-ybhO* (20(G)-200) | *ori*pMB1; [Km$^R$], the HS-*ybhO* (20(G)-200) fragment was amplified by PCR from MG1655 using o6276 and o6416 primers and cloned in the pCR®-BluntII-TOPO® vector. |
| pP177 | pTOPO-*HS-ybhO* (20(G)-60) | *ori*pMB1; [Km$^R$], annealing of complementary o6429 and o6430 primers reconstituting the HS-*ybhO* (20(G)-60) fragment. The annealed primers (generating digested *EcoRI* enzyme restriction site) are cloned in *EcoRI* digested pCR®-BluntII-TOPO® vector. |

| | | |
|---|---|---|
| pP178 | pTOPO-*HS-ybhO* (20(G)-20) | *ori*pMB1; [$Km^R$], annealing of complementary o6431 and o6432 primers reconstituting the HS-*ybhO* (20(G)-20) fragment. The annealed primers (generating digested *EcoRI* enzyme restriction site) are cloned in *EcoRI* digested pCR®-BluntII-TOPO® vector. |
| pP761 | pTOPO-*HS-ybhO* (20(G)-10) | *ori*pMB1; [$Km^R$], annealing of complementary o6478 and o6479 primers reconstituting the HS-*ybhO* (20(G)-10) fragment. The annealed primers (generating digested *EcoRI* enzyme restriction site) are cloned in *EcoRI* digested pCR®-BluntII-TOPO® vector. |
| pP807 | pTOPO-*HS-ybhO* (10(G)-20) | *ori*pMB1; [$Km^R$], annealing of complementary o6472 and o6473 primers reconstituting the HS-*ybhO* (10(G)-20) fragment. The annealed primers (generating digested *EcoRI* enzyme restriction site) are cloned in *EcoRI* digested pCR®-BluntII-TOPO® vector. |
| pP762 | pTOPO-*HS-ybhO* (10(G)-10) | *ori*pMB1; [$Km^R$], annealing of complementary o6480 and o6481 primers reconstituting the HS-*ybhO* (10(G)-10) fragment. The annealed primers (generating digested *EcoRI* enzyme restriction site) are cloned in *EcoRI* digested pCR®-BluntII-TOPO® vector. |
| pQ223 | pTOPO-*HS-ybhO* (9(G)-9) | *ori*pMB1; [$Km^R$], annealing of complementary o6511 and o6512 primers reconstituting the HS-*ybhO* (9(G)-9) fragment. The annealed primers (generating digested *EcoRI* enzyme restriction site) are cloned in *EcoRI* digested pCR®-BluntII-TOPO® vector. |
| pQ342 | pTOPO-*HS-ybhO* (9(G)-8) | *ori*pMB1; [$Km^R$], annealing of complementary o6525 and o6526 primers reconstituting the HS-*ybhO* (9(G)-8) fragment. The annealed primers (generating digested *EcoRI* enzyme restriction site) are cloned in *EcoRI* digested pCR®-BluntII-TOPO® vector. |
| pQ162 | pTOPO-*HS-ybhO* (8(G)-9) | *ori*pMB1; [$Km^R$], annealing of complementary o6513 and o6514 primers reconstituting the HS-*ybhO* (8(G)-9) fragment. The annealed primers (generating digested *EcoRI* enzyme restriction site) are cloned in *EcoRI* digested pCR®-BluntII-TOPO® vector. |
| pQ159 | pTOPO-HS-*ybhO* (10(G)-0) | *ori*pMB1; [$Km^R$], annealing of complementary o6509 and o6510 primers reconstituting the HS-*ybhO* (10(G)-0) fragment. The annealed primers (generating digested *EcoRI* enzyme restriction site) are cloned in *EcoRI* digested pCR®-BluntII-TOPO® vector. |
| pP760 | pTOPO-HS-*ybhO* (0(G)-10) | *ori*pMB1; [$Km^R$], annealing of complementary o6476 and o6477 primers reconstituting the HS-*ybhO* (0(G)-10) fragment. The annealed primers (generating digested *EcoRI* enzyme restriction site) |

| | | |
|---|---|---|
| | | are cloned in *EcoRI* digested pCR®-BluntII-TOPO® vector. |
| pQ706 | pTOPO-*HS-ybhO*-CAGA | *ori*pMB1; [$Km^R$], annealing of complementary o6572 and o6573 primers reconstituting the HS-*ybhO*-CAGA fragment. The annealed primers (generating digested *EcoRI* enzyme restriction site) are cloned in *EcoRI* digested pCR®-BluntII-TOPO® vector. |
| pR102-pR104 | pTOPO-HS-*ybhO*-CCAG | *ori*pMB1; [$Km^R$], the HS-*ybhO*-CCAG fragment was amplified by PCR from pQ706 using o6578 and o6617 primers and cloned in the pCR®-BluntII-TOPO® vector. |
| pR105-pR107 | pTOPO-HS-*ybhO*-TCAG | *ori*pMB1; [$Km^R$], the HS-*ybhO*-TCAG fragment was amplified by PCR from pQ223 using o6578 and o6618 primers and cloned in the pCR®-BluntII-TOPO® vector. |
| pR528 | pTOPO-*HS-alsB* (9(G)-9) | *ori*pMB1; [$Km^R$], annealing of complementary o6648 and o6649 primers reconstituting the HS6 (*9-alsB-9)* fragment. The annealed primers (generating digested *EcoRI* enzyme restriction site) are cloned in *EcoRI* digested pCR®-BluntII-TOPO® vector. |
| p979 | pBAD18 | *ori*ColE1; [$Carb^R$] (Guzman et al. 1995) |
| p3938 | pBAD18*::intI1* | *ori*ColE1; [$Carb^R$] (Demarre et al. 2007) |
| pL290 | pBAD43*::aadA7* | *ori*pSC101; [$Sp^R$] (Vit et al. 2021) |
| pL294 | pBAD43*::aadA7 intI1* | *ori*pSC101; [$Sp^R$] (Vit et al. 2021) |
| pO859-pO860 | pBAD43*::aadA7 intI2**179E | *ori*pSC101; [$Sp^R$], the fragment *intI2* was amplified by PCR from p4327 using o6305 and o6306 primers. The pL294 vector was amplified by PCR using o6307 and o6308 primers. Assembly of these two fragments was achieved by performing Gibson Assembly. |
| pO727-pO728 | pBAD43*::aadA7 intI3* | *ori*pSC101; [$Sp^R$], the fragment *intI3* was amplified by PCR from p4328 using o6309 and o6310 primers. The pL294 vector was amplified by PCR using o6311 and o6312 primers. Assembly of these two fragments was achieved by performing Gibson Assembly. |
| p4327 | pBAD18*::intI2**179E | *ori*ColE1; [$Carb^R$] (Demarre, unpublished ) |
| p4328 | pBAD18*::intI3* | *ori*ColE1; [$Carb^R$] (Demarre, unpublished) |
| pE639 | pLC10 | *oripSC101ts*; [$Sp^R$] Laboratory of David Bikard, Institut Pasteur |
| pM889 | pBAD43-P*tet* | *ori*pSC101; [$Sp^R$], the fragment P*tet* was amplified by PCR from pE639 using o6178 and o6182 primers. The pL294 vector was amplified by PCR using o6181 and o6183 primers. Assembly of these two fragments was achieved by performing Gibson Assembly. |

| | | |
|---|---|---|
| pM888 | pBAD43-P*tet-intI1* | *ori*pSC101; [Sp$^{R}$], the fragment P*tet* was amplified by PCR from pE639 using o6178 and o6179 primers. The pL294 vector was amplified by PCR using o6180 and o6181 primers. Assembly of these two fragments was achieved by performing Gibson Assembly. |
| pN438-pN440 | pBAD43-P*tet*-*ori*pSC101ts | *ori*pSC101*ts;* [Km$^{R}$], amplification by inverse PCR of pE639 using primers o6231 and o6232 |
| pN435-pN437 | pBAD43-P*tet*-intI1-*ori*pSC101ts | *ori*pSC101*ts;* [Km$^{R}$], the fragment *intI1* was amplified by PCR from pL294 using o6234 and o6180 primers. The pE639 vector was amplified by PCR using o6233 and o6179 primers. Assembly of these two fragments was achieved by performing Gibson Assembly. |
| pD060 | pSW23T*::attC*$_{aadA7}$ (bs) | *oriV*$_{\mathrm{R6K}\gamma}$, *oriT*$_{\mathrm{RP4}}$; [Cm$^{R}$] (Nivina et al. 2016) |
| pD059 | pSW23T*::attC*$_{aadA7}$ (ts) | *oriV*$_{\mathrm{R6K}\gamma}$, *oriT*$_{\mathrm{RP4}}$; [Cm$^{R}$] (Nivina et al. 2016) |
| p1880 | pSW23T*::*VCR$_{2/1}$ (bs) | *oriV*$_{\mathrm{R6K}\gamma}$, *oriT*$_{\mathrm{RP4}}$; [Cm$^{R}$] (Biskri et al. 2005) |
| pF486 | pSW23T*::*VCR$_{64}$ (bs) | *oriV*$_{\mathrm{R6K}\gamma}$, *oriT*$_{\mathrm{RP4}}$; [Cm$^{R}$], annealing of complementary and partially overlapping primers (o4329, o4330, o4331, o4332) reconstituting the VCR$_{64}$ site. The annealed primers (generating digested *EcoRI* and *BglI* enzyme restriction sites) are cloned in *EcoRI*/*BglI* digested p1880. |
| p3614 | pSW23T*::attC*$_{dfrB2}$ (bs) | *oriV*$_{\mathrm{R6K}\gamma}$, *oriT*$_{\mathrm{RP4}}$; [Cm$^{R}$] (Demarre, unpublished) |
| p3615 | pSW23T*::attC*$_{ereA2}$ (bs) | *oriV*$_{\mathrm{R6K}\gamma}$, *oriT*$_{\mathrm{RP4}}$; [Cm$^{R}$] (Bouvier et al, 2009) |
| p3616 | pSW23T*::attC*$_{oxa2}$ (bs) | *oriV*$_{\mathrm{R6K}\gamma}$, *oriT*$_{\mathrm{RP4}}$; [Cm$^{R}$] (Bouvier et al, 2009) |
| p2714 | pSW23T*::attI1*(bs) | *oriV*$_{\mathrm{R6K}\gamma}$, *oriT*$_{\mathrm{RP4}}$; [Cm$^{R}$] (Bouvier et al, 2005) |
| p2715 | pSW23T*::attI1*(ts) | *oriV*$_{\mathrm{R6K}\gamma}$, *oriT*$_{\mathrm{RP4}}$; [Cm$^{R}$] (Bouvier et al, 2005) |
| pO323-pO324 | pSW23T*::HS-ybhO* (bs) | *oriV*$_{\mathrm{R6K}\gamma}$, *oriT*$_{\mathrm{RP4}}$; [Cm$^{R}$], *EcoRI*/*BamHI* fragment (*HS-ybhO*) from pO123 cloned in *EcoRI*/*BamHI* digested pDO60. |
| pO321-pO322 | pSW23T*::HS-ybhO* (ts) | *oriV*$_{\mathrm{R6K}\gamma}$, *oriT*$_{\mathrm{RP4}}$; [Cm$^{R}$], *EcoRI*/*BamHI* fragment (*HS-ybhO*) from pO123 cloned in *EcoRI*/*BamHI* digested p2715. |
| pO749-pO750 | pSW23T*::HS-alsB* (bs) | *oriV*$_{\mathrm{R6K}\gamma}$, *oriT*$_{\mathrm{RP4}}$; [Cm$^{R}$], *EcoRI*/*BamHI* fragment (*HS-aslB*) from pO127 cloned in *EcoRI*/*BamHI* digested pDO60. |
| pO751 | pSW23T*::HS-alsB* (ts) | *oriV*$_{\mathrm{R6K}\gamma}$, *oriT*$_{\mathrm{RP4}}$; [Cm$^{R}$], *EcoRI*/*BamHI* fragment (*HS-aslB*) from pO127 cloned in *EcoRI*/*BamHI* digested p2715. |
| pO752 | pSW23T*::HS-pyrE* (bs) | *oriV*$_{\mathrm{R6K}\gamma}$, *oriT*$_{\mathrm{RP4}}$; [Cm$^{R}$], *EcoRI*/*BamHI* fragment (*HS-pyrE*) from pO125 cloned in *EcoRI*/*BamHI* digested pDO60. |
| pO753-755 | pSW23T*::HS-pyrE* (ts) | *oriV*$_{\mathrm{R6K}\gamma}$, *oriT*$_{\mathrm{RP4}}$; [Cm$^{R}$], *EcoRI*/*BamHI* fragment (*HS-pyrE*) from pO125 cloned in *EcoRI*/*BamHI* digested p2715. |
| p7848 (pMP7) | pSW23T*::araC* P$_{\mathrm{BAD}}$-*ccdB* | *oriV*$_{\mathrm{R6K}\gamma}$, *oriT*$_{\mathrm{RP4}}$; [Cm$^{R}$] (Val et al. 2012) |

| | | |
|---|---|---|
| pM779-pM781 | pMP7::*attC*$_{aadA7}$ | *oriV*$_{R6K\gamma}$, *oriT*$_{RP4}$; [Cm$^{R}$], annealing of complementary o6075 and o6076 primers reconstituting the *attC*$_{aadA7}$ site. The annealed primers (generating digested *EcoRI* and *BamHI* enzyme restriction sites) are cloned in *EcoRI*/*BamHI* digested p7848. |
| pN695-pN697 | pSW23T*::attC*$_{aadA7}$*-km* | *oriV*$_{R6K\gamma}$, *oriT*$_{RP4}$; [Cm$^{R}$], the *km* CDS was amplified by PCR from the p929 plasmid using o6197 and o6198 primers. This *BamHI*/*SpeI* digested fragment was cloned in the *BamHI*/*SpeI* digested pD060 plasmid. |
| pN708-pN709 | pSW23T*::attC*$_{aadA7}$*-RBS1ggagg-km* | *oriV*$_{R6K\gamma}$, *oriT*$_{RP4}$; [Cm$^{R}$], the *attC*-RBS1 fragment was amplified by PCR using SWbeg and o6262 primers and the RBS1-*km* fragment using o6261 and SWend primers. Both PCR were performed using the pN696 plasmid as template. These two fragments were assembled by PCR using SWbeg and SWend primers. The obtained fragment was then digested by *EcoR/SpeI* and cloned in the *EcoRI*/*SpeI* digested pD060 plasmid. |
| pN705-pN707 | pSW23T*::attC*$_{aadA7}$*-RBS2aggag-km* | *oriV*$_{R6K\gamma}$, *oriT*$_{RP4}$; [Cm$^{R}$], the *attC*-RBS2 fragment was amplified by PCR using SWbeg and o6260 primers and the RBS2-*km* fragment using o6259 and SWend primers. Both PCR were performed using the pN696 plasmid as template. These two fragments were assembled by PCR using SWbeg and SWend primers. The obtained fragment was then digested by *EcoR/SpeI* and cloned in the *EcoRI*/*SpeI* digested pD060 plasmid. |
| p8701 | pMP96 | *ori*pSC101ts; [Sp$^{R}$] (Val et al. 2012) |
| pM335 | pMP96Δint-xis-oriT | *ori*pSC101ts; [Sp$^{R}$], amplification by inverse PCR of p8701 using o6038 and o6039 primers. |

| **Primers** | **Sequences** |
|---|---|
| ***Primers used for plasmid construction*** | |
| o3849 | AATTCGTCTAACAATTCATTCAAGCCGACGCCGCTTCGCGGCGCGGCTTAATTCAAGCGTTAGACG |
| o3850 | GATCCGTCTAACGCTTGAATTAAGCCGCGCCGCGAAGCGGCGTCGGCTTGAATGAATTGTTAGACG |
| o4329 | AATTCAGATCTGGTTATAACAAACGCC |
| o4330 | TTAAGAGGGACTGCCAACGCGTGGCATTTCCAGTCCCAATGAGCCGTGGTGGTTACGGTTGTTGTGTTTGAGTTTCGTTGTTATGCGTTGTCAGCCCCTT |
| o4331 | GGGCTGACAACGCATAACAACGAAACTCAAACACAACAACCG |
| o4332 | TAACCACCACGGCTCATTGGGACTGGAAATGCCACGCGTTGGCAGTCCCTCTTAAGGCGTTTGTTATAACCAGATCTG |
| o4870 | AATTCGCATAACCTGCCAATCCACCGGACGGTTTTCAACCGCCGGTGATCAGCGCGTTATGCG |

| | |
|---|---|
| o4871 | GATCCGCATAACGCGCTGATCACCGGCGGTTGAAAACCGTCCGGTGGATTGGCAGGTTATGCG |
| o6035 | AATGATACGGCGACCACCGAGATCTACACGATTAGCAACACTCTTTCCCTACACGACGCTCTTCCGATCTCAATTCATTCAAGCCGACGCCGCTTCGCG |
| o6036 | CAAGCAGAAGACGGCATACGAGAT |
| o6038 | TAATACTAGTAGCGGCCGCTGC |
| o6039 | CTAGAGGATCCCCGGGTACC |
| o6075 | AATTCGTCTAACAATTCATTCAAGCCGACGCCGCTTCGCGGCGCGGCTTAATTCAAGCGTTAGACCTGCA |
| o6076 | GGTCTAACGCTTGAATTAAGCCGCGCCGCGAAGCGGCGTCGGCTTGAATGAATTGTTAGACG |
| o6178 | CCCTATGCTACTCCGTCAAGCCGTCAATTGTCTGATTCGTTACCAACACGCCTGCCAGGAATTGGGGATCGGTTAAGACCC |
| o6179 | CCTTGACCGAACGCAGCGGTGGTAACGGCGCAGTGGCGGTTTTCATTTTTGCCTCCTAACTAGGTCATTTGATATGCCTCCGG |
| o6180 | CCGGAGGCATATCAAATGACCTAGTTAGGAGGCAAAAATGAAAACCGCCACTGCGCCGTTACCACCGCTGCGTTCGGTCAAGG |
| o6181 | GGGTCTTAACCGATCCCCAATTCCTGGCAGGCGTGTTGGTAACGAATCAGACAATTGACGGCTTGACGGAGTAGCATAGGG |
| o6182 | CGCAGGGGTAGTGAATCCGCCAGGATTGACTTGCGCTGCCTTTTGCCTCCTAACTAGGTCATTTGATATGCCTCCGG |
| o6183 | CCGGAGGCATATCAAATGACCTAGTTAGGAGGCAAAAGGCAGCGCAAGTCA<br>ATCCTGGCGGATTCACTACCCCTGCG |
| o6197 | GGCCGGGATCCGAGAACTATGATTGAACAAGATGGATTGCACGC |
| o6198 | GCGGACTAGTTCAGAAGAACTCGTCAAGAAGGCGATAGAAGGC |
| o6231 | ATTCACCACCCTGAATTGACTCTCTTCCGGGCGCTATCATGCC |
| o6232 | TTTTGCCTCCTAACTAGGTCATTTGATATGCCTCCG |
| o6233 | CGCTTGATGCGCTGCCGCCCCTCACTAGTGAGAGGTAGATTCACCACCCTGAATTGACTCTCTTCCG |
| o6234 | CGGAAGAGAGTCAATTCAGGGTGGTGAATCTACCTCTCACTAGTGAGGGGCGGCAGCGCATCAAGCG |
| o6259 | CGCGGCTTAATTCAAGCGTTAGACAGGAGCGAGAACTATGATTGAACAAGATGGATTGCA |
| o6260 | TGCAATCCATCTTGTTCAATCATAGTTCTCGCTCCTGTCTAACGCTTGAATTAAGCCGCG |
| o6261 | CGCGGCTTAATTCAAGCGTTAGACGGAGGCGAGAACTATGATTGAACAAGATGGATTGCA |
| o6262 | TGCAATCCATCTTGTTCAATCATAGTTCTCGCCTCCGTCTAACGCTTGAATTAAGCCGCG |
| o6273 | GCCGAATTCGGTGGTAAGCAATATGTCTCTACAGG |
| o6274 | GCCGGATCCAAAGTCGCACCACGAAATCTTGACGACTATTACC |
| o6275 | GCCGAATTCTGGCGTAAAAAGTGGAACGCCAGC |
| o6276 | GCCGGATCCGATGATATTGAACGCCATTATTTGAAAATGC |
| o6277 | GCCGGATCCCGTCAACATCATGAAAGCAGTCTGAAAGTGG |
| o6278 | GCCGAATTCCTATTGTACTCCTGAATCGTCCGGGACG |
| o6279 | GCCGGATCCATGCCGGAATCCACCAACGCTTCAGC |
| o6280 | GCCGAATTCAAGGGATGGTAGCCGCAGTTTCTGTCG |
| o6281 | GCCGGATCCGCGCAGACCCGAAAGTATCTGACTCC |
| o6282 | GCCGAATTCGCCGAGAATACCGATAACACCACC |
| o6283 | GCCGGATCCTGCGCCCCGAGGAGTTCGGTGAGTTTCTGG |

| | |
|---|---|
| o6284 | GCCGAATTCGGCCGGGCGACCGTGCGCCAGC |
| o6293 | GCCGGATCCTTCGCGTTGTAGAGTGAATTCATCAGG |
| o6294 | GCCGAATTCCCGGTTGCGACATATTCAACATTCG |
| o6299 | GCCGAATTCTACTGGCCGATTATGTCTTCTATTCCG |
| o6300 | GCCGGATCCTTAATCGCTGCCCCCAGGCTTGC |
| o6301 | GCCGAATTCTGCCGTTCAGATTATCGCGCAGCG |
| o6303 | GCCGAATTCCTGAGCGGATCGAGATTACTGGACCC |
| o6304 | GCCGAATTCGATCGTCCATCAATGCCACTTTGCC |
| o6305 | GCAGGAGGAATTCGAGCTCTAACAAAGGAGCAAGCCATGTCTAACAGTC CATTTTTAAATTCTATACGCACGGATATGCG |
| o6306 | GGGTAGTGAATCCGCCAGGATTGACTTGCGCTGCCTTTACTGATTGATAA GTAGCATCAGTCCATCCGCAGGA |
| o6307 | CGCATATCCGTGCGTATAGAATTTAAAAATGGACTGTTAGACATGGCTTG CTCCTTTGTTAGAGCTCGAATTCCTCCTGC |
| o6308 | TCCTGCGGATGGACTGATGCTACTTATCAATCAGTAAAGGCAGCGCAAGT CAATCCTGGCGGATTCACTACCC |
| o6309 | GCAGGAGGAATTCGAGCTCTAACAAAGGAGCAAGCCATGAACAGGTATA ACGGATCTGCCAAACCTGACTGGGTCCC |
| o6310 | GGGTAGTGAATCCGCCAGGATTGACTTGCGCTGCCTCAGCCGGGCGACA AGTGCAAGGCCAAGGCGTCC |
| o6311 | GGACGCCTTGGCCTTGCACTTGTCGCCCGGCTGAGGCAGCGCAAGTCAAT CCTGGCGGATTCACTACCC |
| o6312 | GGGACCCAGTCAGGTTTGGCAGATCCGTTATACCTGTTCATGGCTTGCTC CTTTGTTAGAGCTCGAATTCCTCCTGC |
| o6313 | GCCGAATTCATGACATTTGCTTCGAGATTCAGT |
| o6314 | AATTCCAAGTGATTATATTAATTAACGGTAAGCATCAGCGGGTGACAAAA CG AGCATGCTTACTAATAAAATGTTAACCTCTGAGGAAGG |
| o6315 | GATCCCTTCCTCAGAGGTTAACATTTTATTAGTAAGCATGCTCGTTTTGTC ACCCGCTGATGCTTACCGTTAATTAATATAATCACTTGG |
| o6316 | AATTCGGTGCCGTGCGACTTTGTTTAACGACCACGGTTGTGGGTATCCGG TGTTTGGTCAGATAAACCACAAGTTAGATGCACTAAGCAG |
| o6317 | GATCCtGCTTAGTGCATCTAACTTGTGGTTTATCTGACCAAACACCGGATA CCCACAACCGTGGTCGTTAAACAAAGTCGCACGGCACCG |
| o6414 | GCCGAATTCTTGCCGTGGAGCGGGCGGCGGCGGTACT |
| o6415 | GCCGAATTCGCGGTACTCAAAAACCTGAACGCC |
| o6416 | GCCGAATTCCGCCGCCTTTAACCAGATAGTTATACAGC |
| o6429 | AATTCGCCCTTGCCGGATCCGATCATTCAGGGCGAACCGGATATGCCGAT TGTCAGAGTCGGTGCGCGCTTGCTGTATAACTATCTGGTTAAAGGCGGCG GAATTCGGAAGGGCG |
| o6430 | AATTCGCCCTTCCGAATTCCGCCGCCTTTAACCAGATAGTTATACAGCAA GCGCGCACCGACTCTGACAATCGGCATATCCGGTTCGCCCTGAATGATCG GATCCGGCAAGGGCG |
| o6431 | AATTCGCCCTTGCCGGATCCGGTGCGCGCTTGCTGTATAACTATCTGGTTA AAGGCGGCGGAATTCGGAAGGGCG |
| o6432 | AATTCGCCCTTCCGAATTCCGCCGCCTTTAACCAGATAGTTATACAGCAA GCGCGCACCGGATCCGGCAAGGGCG |
| o6472 | AATTCGCCCTTGCCGGATCCGGTGCGCGCTTGCTGTATAACTATCTGGTTG AATTCGGAAGGGCG |
| o6473 | AATTCGCCCTTCCGAATTCAACCAGATAGTTATACAGCAAGCGCGCACCG GATCCGGCAAGGGCG |
| o6476 | AATTCGCCCTTGCCGGATCCTGCTGTATAACGAATTCGGAAGGGCG |
| o6477 | AATTCGCCCTTCCGAATTCGTTATACAGCAGGATCCGGCAAGGGCG |
| o6478 | AATTCGCCCTTGCCGGATCCTGCTGTATAACTATCTGGTTAAAGGCGGCG GAATTCGGAAGGGCG |
| o6479 | AATTCGCCCTTCCGAATTCCGCCGCCTTTAACCAGATAGTTATACAGCAG GATCCGGCAAGGGCG |

| | |
|---|---|
| o6480 | AATTCGCCCTTGCCGGATCCTGCTGTATAACTATCTGGTTGAATTCGGAA GGGCG |
| o6481 | AATTCGCCCTTCCGAATTCAACCAGATAGTTATACAGCAGGATCCGGCAA GGGCG |
| o6509 | AATTCGCCCTTGCCGGATCCTAACTATCTGGTTGAATTCGGAAGGGCG |
| o6510 | AATTCGCCCTTCCGAATTCAACCAGATAGTTAGGATCCGGCAAGGGCG |
| o6511 | AATTCGCCCTTGCCGGATCCGCTGTATAACTATCTGGTGAATTCGGAAGG GCG |
| o6512 | AATTCGCCCTTCCGAATTCACCAGATAGTTATACAGCGGATCCGGCAAGG GCG |
| o6525 | AATTCGCCCTTGCCGGATCCCTGTATAACTATCTGGTGAATTCGGAAGGG CG |
| o6526 | AATTCGCCCTTCCGAATTCACCAGATAGTTATACAGGGATCCGGCAAGGG CG |
| o6513 | AATTCGCCCTTGCCGGATCCGCTGTATAACTATCTGGGAATTCGGAAGGG CG |
| o6514 | AATTCGCCCTTCCGAATTCCCAGATAGTTATACAGCGGATCCGGCAAGGG CG |
| o6572 | AATTCGCCCTTGCCGGATCCTCTGTATAACTATCTGGTGAATTCGGAAGG GCG |
| o6573 | AATTCGCCCTTCCGAATTCACCAGATAGTTATACAGAGGATCCGGCAAGG GCG |
| o6578 | GGATCCGGCAAGGGCGAATTCTGCAGATATCCA |
| o6593 | GGATCCTCTAGAGTCGACCTGCAGGC |
| o6594 | GATGCCTAACTTTGTTTTAGGGC |
| o6605 | CCCTAAAACAAAGTTAGGCCAGGGATCCTCTAGAGTCGACC |
| o6606 | GGTCGACTCTAGAGGATCCCTGGCCTAACTTTGTTTTAGGG |
| o6607 | CCCTAAAACAAAGTTAGGTCAGGGATCCTCTAGAGTCGACC |
| o6608 | GGTCGACTCTAGAGGATCCCTGACCTAACTTTGTTTTAGGG |
| o6617 | CTGGTATAACTATCTGGTGAATTCGGAAGGG |
| o6618 | CTGATATAACTATCTGGTGAATTCGGAAGGG |
| o6619 | GCGGATCCTCTAGAGTCGACCTGCAGGC |
| o6620 | TGCCTAACTTTGTTTTAGGGCGACTGCC |
| o6648 | AATTCGCCCTTGCCGGATCCGGCCTGTGATCACTATGTTGAATTCGGAAG GGCG |
| o6649 | AATTCGCCCTTCCGAATTCAACATAGTGATCACAGGCCGGATCCGGCAAG GGCG |
| o6699 | GCCGAATTCGGCTTGTTATGACTGTTTTTTTGTACAG |
| o6700 | GCCGGATCCGATGCCTAACTTTGTTTAGGGCGACTGC |
| o6716 | AATTCGCCCTTGCCGGATCCGCATAACGCGCTGATCACCGGCGGTTGAAA ACCGTCCGGTGGATTGGCAGGTTATGCGAATTCGGAAGGGCG |
| o6717 | AATTCGCCCTTCCGAATTCGCATAACCTGCCAATCCACCGGACGGTTTTC AACCGCCGGTGATCAGCGCGTTATGCGGATCCGGCAAGGGCG |
| ***Primers used to determine the insertion/excision cassette site*** | |
| SWbeg | CCGTCACAGGTATTTATTCGGCG |
| SWend | CCTCACTAAAGGGAACAAAAGCTG |
| MRV | AGCGGATAACAATTTCACACAGGA |
| MFD | CGCCAGGGTTTTCCCAGTCAC |
| o1366 | AGCGGGTGTTCCTTCTTCACTG |
| o1388 | CCGGGCAGGATAGGTGAAGTAG |
| o1704 | AGAGAACATAGCGTTGCCTTGG |
| o1714 | TTTGTCAGCAAGATAGCCAG |
| o1863 | GGCCACGCGTCGACTAGTACNNNNNNNNNNACGCC |
| o1865 | GGCCACGCGTCGACTAGTAC |

| | |
|---|---|
| o2405 | ATACGACTCACTATAGGGCG |
| o6263 | GCTTGTTACCGTCATTATCATCCG |
| o6264 | GGCGCTGAAGGTGGTGAAGTCGC |
| o6265 | CCAGCAGCTCGCGAACCTCGG |
| o6266 | CCTGGCAGATGGAAGGTGCC |
| o6267 | GGTTGATGCGATGATCCAGGGC |
| o6268 | GCTGTGGAACCTTGCGGGAAAGC |
| ***Primers used to perform the digital PCR*** | |
| oriC_MG1655_F | CCACCGAGAAGAACATGGAG |
| oriC_MG1655 55_5 | GCCGCAGGATTACATAGGAC |
| ori C probe | ATTGTCCAGAAGGTGGCTGGGGGGTTTT |
| terC_MG1655_F | TTTCGTAACCGACAGCATAGG |
| terC_MG1655 55_5 | GTCCACTCCGGTATCAGAAATG |
| ter C probe | TTCAGCATCTGTCGCCTGACTTCA |
